# The RNA-Binding Protein DND_1_ Acts Sequentially as a Negative Regulator of Pluripotency and a Positive Regulator of Epigenetic Modifiers Required for Germ Cell Reprogramming

**DOI:** 10.1101/402008

**Authors:** Victor A. Ruthig, Matthew B. Friedersdorf, Jason A. Garness, Steve C. Munger, Corey Bunce, Jack D. Keene, Blanche Capel

## Abstract

The adult spermatogonial stem cell population arises from pluripotent primordial germ cells (PGCs) that enter the fetal testis around embryonic day 10.5 (E10.5). These cells undergo rapid mitotic proliferation, then enter a prolonged period of cell cycle arrest (G1/G0) during which they transition to pro-spermatogonia. In mice homozygous for the *Ter* mutation in the RNA-binding protein *DND_1_* (*DND_1_*^*Ter/Ter*^), many germ cells fail to enter G1/G0, and give rise to teratomas, tumors in which many embryonic cell types are represented. To investigate the origin of these tumors, we sequenced the transcriptome of male germ cells in *DND_1_*^*Ter/Ter*^ mutants at E_12.5_, E_13.5_, and E_14.5_, just prior to the formation of teratomas, and correlated this information with direct targets of DND_1_ identified by DO-RIP-Seq. Consistent with previous results, we found that DND_1_ controls the down regulation of many genes associated with pluripotency and active cell cycle, including elements of the mTor, Hippo and Bmp/Nodal signaling pathways. However, DND_1_ targets also include genes associated with male differentiation including a large group of chromatin regulators activated in wild type but not mutant germ cells during the transition between E_13.5_ and E_14.5_. These results suggest multiple functions of DND_1_, and link DND_1_ to the initiation of epigenetic modifications in male germ cells.

## Introduction

Primordial germ cells (PGCs) are specified at the base of the allantois in mouse embryos at embryonic day (E)6.75, proliferate, migrate through the gut mesentery, and arrive in the gonad between E10.0-E11.0, expressing stem cell markers including SOX2 (SRY-box 2), NANOG (Nanog homeobox), and OCT4 (also known as POU5F1, POU domain, class 5, transcription factor 1) (Matsui et al. (1992); (McLaren, 1995); (Yeom et al., 1996); (Pesce and Scholer, 2000; Yamaguchi et al., 2005). Prior to E_12.5_, germ line stem cells (EGCs) can be derived readily from the germ cell population (Matsui et al., 1992), and teratomas, a tumor in which all embryonic cell types are represented, spontaneously arise from germ cells in males with a frequency of 1-10% in some strains (Stevens and Little, 1954); (Bustamante-Marin et al., 2013); (Dawson et al., 2018). However, between E_12.5_ and E_15.5_, the efficiency of both EGC derivation and teratoma induction declines, presumably reflecting changes that lead to suppression of the underlying pluripotent state of germ cells. These changes are coincident with the entry of male germ cells into G1/G0 cell cycle arrest and their fate transition to pro-spermatogonia (Fig. 1A) (Stevens, 1966); (Western et al., 2011; Western et al., 2010).

**Figure 1.**
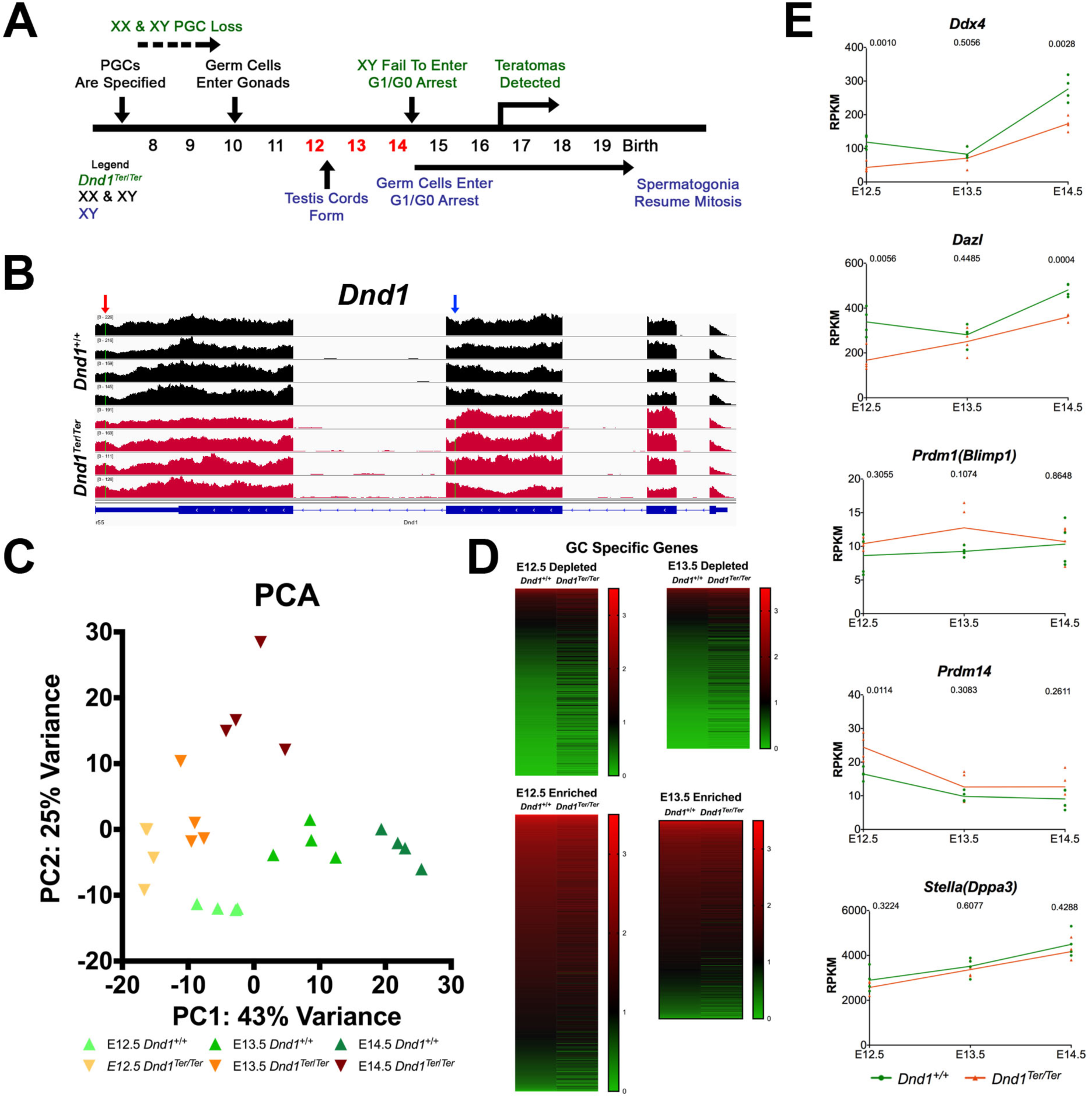
Male germ cells in *DND_1_*^*Ter/Ter*^ mutants retain a germ cell identity. **(A)** Time line of male germ cell development and defects in *DND_1_*^*Ter/Ter*^ mutants. Primordial germ cells (PGCs) are specified at ~E7.5, migrate and arrive in the gonad at E10.5. In the male gonad, they are enclosed inside testis cords by E_12.5_, and undergo G0 arrest between E_14.5_-E_15.5_, remaining in arrest until after birth. In *DND_1_*^*Ter/Ter*^ mutants, many PGCs are lost soon after specification. Remaining germ cells enter the gonad, but fail to arrest in G0. Teratomas are detected in the fetal testis beginning at ~E16.5 (shown in green above time line). **(B)** *DND_1_* is transcribed in mutants similar to wild type levels at E_12.5_. The *Ter* mutation is visible in the 2^nd^ exon of all mutant samples (blue arrow, red samples), whereas the 129T2-specific SNP (red arrow) is present in all samples in the 5, non-coding region. **(C)** A PCA analysis revealed that mutant germ cells are more similar to wild type at E_12.5_ and E_13.5_ and diverge further at E_14.5_. **(D)** Nonetheless, male germ cells show an overall expression pattern characteristic of germ cells with respect to genes that are normally depleted (top) and genes that are normally up-regulated (bottom) at E_12.5_ and E_13.5_. **(E)**. Specific examples of gene expression across the timecourse (green line = wild type samples; orange line = *DND_1_*^*Ter/Ter*^ in all graphs in the manuscript; significance values for each time point are shown at top of graph). While both *Ddx4* (*Vasa*) and *Dazl* show a delay in activation, similar levels at E_13.5_, and a slight down regulation at E_14.5_, *Prdm1* (*Blimp1*), *Prdm14*, and *Stella (Dppa3)* show no significant differences from wild type at any stage.

A mutation, called *Ter*, that arose spontaneously during routine breeding of 129/SvJ mice, caused a severe reduction in germ cell numbers soon after their specification, and acted as a strong modifier that increased the incidence of spontaneous testicular teratomas to >90% in 129/SvJ mutants (Fig. 1A)(Stevens, 1973). In 2005, *Ter* was mapped to a point mutation that introduced a premature stop codon in the RNA-binding protein (RBP) Dead End 1 *DND_1_* (*DND_1_*^*Ter*^) (Fig. 1B) (Youngren et al., 2005). Understanding how DND_1_ is involved in stabilization of germ cell fate will provide important insights into how this critical process is effected in germ cells and, by analogy, in other stem cell populations from which tumors arise.

The function of DND_1_ has been studied by knockouts in zebrafish (Gross-Thebing et al., 2017; Weidinger et al., 2003) and mouse (Zechel et al., 2013), and by a conditional deletion at E_13.5_ in mouse (Suzuki et al., 2016), but none of these mutants formed teratomas, indicating that there is something unique about the *DND_1_*^*Ter*^ mutation and its 129/SvJ genetic background. We have focused our efforts on characterizing the transcriptome in germ cells from *DND_1_*^*Tet/Ter*^ 129SvT2/SvEMSJ (129T2) mutant embryos just prior to the time when teratomas form (Cook et al., 2009; Cook et al., 2011; Heaney et al., 2012), and cross-referencing this information with direct binding targets of DND_1_.

Previous studies have implicated DND_1_ in both positive and negative regulatory roles. The molecular role of DND_1_ was first investigated in a human tumor cell line where the protein was shown to bind to the mRNAs of *Cdkn1b* (a negative regulator of the cell cycle) and *Lats2* (a tumor suppressor and negative regulator of p53) to protect these transcripts from miRNA-mediated translational repression (Kedde et al., 2007) perhaps through interaction with APOBEC3 (apolipoprotein B mRNA editing enzyme, catalytic polypeptide-like) (Bhattacharya et al., 2008). Other studies identified targets protected by DND_1_ in NIH_3_T_3_ or HEK_293_ cells, including the negative cell cycle regulator *Cdkn1b* (cyclin dependent kinase inhibitor 1B)(Cook et al., 2011) and *Ezh2* (enhancer of zeste homolog 2), a mediator of H_3_K_27_me_3_ repression (Gu et al., 2018). Subsequently, two labs showed that DND_1_ acts as a negative regulator of mRNAs by recruiting targets to the CCR4-NOT deadenylase complex for degradation (Suzuki et al., 2016); (Yamaji et al., 2017). Suzuki et al. showed that DND_1_ acts as an essential partner of NANOS_2_ (C_2_HC-type zinc finger 2, another male-specific RBP) to form P-bodies, and to select and recruit mRNA for degradation. In these studies, conditional deletion of *DND_1_* at E_13.5_ led to up-regulation of proliferation, meiosis markers, and Caspase_3_, and the down regulation of several NANOS_2_ targets associated with male differentiation, similar to *NANOS_2_* mutants (Suzuki et al., 2016).

Early studies relied on a candidate approach to identify targets of DND_1_, which included pluripotency factors and cell cycle genes (Cook et al., 2011; Kedde et al., 2007; Zhu et al., 2011). More recently, Yamaji and co-workers performed photoactivatable-Ribonuleoside-Enhanced Cross-Linking and Immunoprecipitation (PAR-CLIP) assays in HEK_293_ cells, and investigated the expression of identified targets in induced primordial germ like cells (PGLCS) and germ line stem cells derived from adult spermatogonia. Consistent with Suzuki,s results, Yamaji et al. showed that DND_1_ acts with CCR_4_-NOT to destabilize and repress RNAs associated with apoptosis, inflammation and signaling pathways including TgfB, Wnt, and PI3K-AKT. Extrapolating results from these in vitro cell types, Yamaji et al. suggested that activation of pluripotency genes, as well as TgfB and inflammatory signaling genes, contribute to the formation of teratomas in gonadal germ cells (Yamaji et al., 2017).

We sequenced the transcriptome of male germ cells in *DND_1_*^*Ter/Ter*^ mutants at E_12.5_, E_13.5_, and E_14.5_, just prior to the formation of teratomas, and correlated this information with direct targets of DND_1_. We identified direct transcript targets of DND_1_ using an independent method, Digestion Optimized-RIP-seq (DO-RIP-seq), after expressing a tagged DND_1_ protein in HEK_293_ cells. We cross-referenced our DO-RIP-seq targets with the PAR-CLIP results from Yamaji et al., and compared the transcript levels of nontargets and targets common to both RNA-immunoprecipitation assays in isolated germ cells from 129T2-*DND_1_*^*Ter/Ter*^ and *DND_1_*^+/+^ littermates at E_12.5_, E_13.5_ and E_14.5_. Consistent with our previous results and the molecular models proposed by Suzuki and Yamaji, between E_12.5_-E_13.5_ DND_1_ controls the down regulation of many genes associated with pluripotency and cell cycle, including elements of the both the Hippo and Bmp/Nodal signaling pathways. However, we found that DND_1_targets include many genes associated with male differentiation including a large group of chromatin regulators normally activated during the second transition between E_13.5_ and E_14.5_. These results suggest multiple functions of DND_1_, and link DND_1_ to the initiation of epigenetic modifications in male germ cells.

## Materials and Methods

### Mice, timed matings, and genotyping

*DND_1_*^*Ter/+*^ mice were kindly provided by Dr. Joseph Nadeau and maintained on a 129T2/SvEmsJ background. 129T2/SvEmsJ (hereafter referred to as “129T2”; www.jax.org/strain/002065) was predicted to be the closest living strain at The Jackson Laboratory (JAX) to the original 129/SvJ strain on which *Ter* arose. Because the formation of teratomas is a 129-strain-specific phenotype, to generate mutants in which germ cells were fluorescently tagged, Oct4-EGFP (originally on a mixed genetic background) (Yoshimizu et al., 1999) was back-crossed to 129T2-*DND_1_*^*Ter/+*^ for 9 generations, then crossed to 129T2-*DND_1_*^*Ter/Ter*^ mutant females to generate 129T2-*DND_1_*^*Ter/+*^;*Oct4-EGFP* male and female heterozygotes. For timed matings, heterozygotes were intercrossed, females were checked for plugs, and staged as day E0.5 if positive. For genotyping, tail DNA was extracted using standard methods and genotyped using the following primer sets: GFP-F 5-AAG TTC ATC TGC ACC ACC G and GFP-R 5-TCC TTG AAG AAG ATG GTG CG and Ter-F 5-GTA GTT CAG GAA CTC CAC TTG TG-3, Ter-R 5-GCT CAA GTT CAG TAC GCA C-3. *DND_1_*^*Ter*^ mice were genotyped by PCR using an annealing temperature of 62°C. The PCR product (145 bp) was digested overnight at 37°C with the restriction enzyme *DdeI* and run on a 4% agarose gel or 6% acrylamide gel. *DdeI* digestion of DNA from mice with the *DND_1_*^*Ter*^ mutation produces 123 bp and 22 bp products.

### FACS, RNA isolation, Library Preparation

Global gene expression was quantified by RNA sequencing (RNA-seq) in germ cells at multiple embryonic stages. To isolate germ cells, embryos were collected at E_12.5_, E_13.5_, and E_14.5_. The gonad was isolated from XY embryos, and incubated in 250 ml 0.25% Trypsin EDTA (Gibco #25200) at 37**^o^**C for 5–10 minutes. The trypsin was removed and replaced with 400 ml PBS with or without 4 ml RNase-free DNase (Promega **#**M6101). The tissue was dissociated, and the cells were passed through a 20μm cell strainer (BD Falcon **#**352235). Fluorescence activated cell sorting (FACS) was performed by the Duke Comprehensive Cancer Center Flow Cytometry Shared Resource. Between 200-900 EGFP+ cells were sorted into 100 μl Qiagen Buffer RLT (from RNeasy Micro Kit, Qiagen #74004) +DTT, without carrier RNA and frozen at **-**80**^o^**C.

Total RNA was extracted from purified germ cell aliquots and eluted in 13 μl of RNase/DNase free water using the Qiagen RNeasy Micro Kit (Qiagen #74004) with on column DNase digestion following manufacturer,s protocols. Next, 10 μl of RNA eluate went directly into first strand synthesis, using oligo-dT primers according to the Smart-seq2 protocol (Picelli et al., 2014) scaled for input. Pre-amplification mix was prepared according to the Smart-seq2 protocol, scaled to 27 μl input from the first strand synthesis reaction. DNA was purified using the recommended Agencourt AMPure XP bead kit (Beckman Coulter #A63880), DNA was quantified by fluorometric quantitation (QUBIT). 0.5 ng from each sample was brought forward for tagmentation using the Nextera XT DNA Sample Prep Kit (illumina #15032352). Samples were size-selected using a 0.6 ratio of SPRIselect beads (Beckman Coulter #B23317) to DNA to obtain fragments >300 bp. Samples were eluted from beads according to protocol, and submitted to the Duke Sequencing Facility for QC by QUBIT and by traces run on an Agilent 2100 bioanalyzer to determine library sizes. The concentration of each library was normalized and 8 pmoles of the pool were sequenced using 50bp SR on the Illumina Hi-seq 2500. Eight samples (4 mutant and 4 wild type) were pooled for each stage and run in technical replicate in 2 lanes of a flow cell. The Duke Sequencing Facility used FastQC for preliminary quality control.

### Transcriptome processing and mapping reads

There were ~22M reads/sample at E_12.5_, ~15M reads/sample at 13.5, and ~38M reads/sample at 14.5. Bowtie 2 was used to align the RNA-seq reads to mm9 using default settings. >81% of E_12.5_ transcripts aligned to the genome, whereas that number for >87% at E_13.5_, and >92% at E_14.5_. The Cufflinks pipeline (version 2.2.0) (Top-Hat, Cufflinks, Cuffmerge) was used to assemble transcripts using default settings. We considered only those transcripts previously annotated using Ensembl75. Gene levels were quantified using Cuffquant, and differential expression analysis was performed using Cuffdiff. This data was used for further analyses.

**Filtering the datasets** We found that some of the most significant outlier genes are strongly expressed in Leydig and Sertoli cells based on the Jameson datasets (Jameson et al., 2012) (Suppl. Fig. 1). Most of these genes were upregulated between E_13.5_ and E_14.5_ in mutant samples. We considered the possibility that this might reflect partial transdifferentiation of germ cells to express genes associated with somatic lineages, but using fluorescent immunocytochemistry we did not detect any co-localization of markers (data not shown). Because the intensity of *Oct4-EGFP* fluorescence declines, and the number of germ cells in mutants is reduced at E_14.5_, we suspected that contamination by somatic cell types in the gonad during fluorescence activated cell-sorting (FACS) accounted for this problem. As a means to eliminate any transcripts that might be due to contamination by another cell type, cell-type-specific microarray data from the Jameson et al. study, for stages E11.5, E_12.5_ and E_13.5_, was used to generate lists of all genes enriched in the male germ cell population. Lists from E11.5 and E_12.5_ in the Jameson array were applied to the E_12.5_ and E_13.5_ transcriptome data sets. The list from E_13.5_ in the Jameson array was applied to the E_13.5_ and E_14.5_ transcriptome data sets. Only genes appearing on these lists were retained in the filtered RNA-seq data (1213 genes for E_12.5_-E_13.5_, and 1207 genes for E_13.5_-E_14.5_). The filtered datasets were replotted and used for this analysis. A similar method was used to generate lists of genes specifically enriched and depleted in all germ cells, female germ cells and male germ cells at E_12.5_ and E_13.5_ (E_12.5_ or E_13.5_ lists from Jameson Array applied to E_12.5_ or E_13.5_ transcriptome data sets respectively). These lists were used for heat map analysis.

**RepEnrich Analysis** To evaluate the status of repetitive elements (specifically transposable elements), the original unfiltered raw data for wild type and *DND_1_*^*Ter*/*Ter*^ samples were compared using RepEnrich (Criscione et al., 2014). The protocol for library preparation, mapping and differential enrichment analysis (Robinson et al., 2010) was followed as described by (Greco et al., 2016) and detailed by SW Criscione (https://github.com/nskvir/RepEnrich) with minor adjustments. Briefly, Bowtie 1.2.1 was used (Langmead et al., 2009) and the custom repetitive annotation was prepared using parameters: -p 6 -q -m 1 -S --max. For statistical analysis, a FDR (false discovery rate) of less than 0.05 was used as the cutoff for significance (Greco et al., 2016); (Criscione et al., 2014).

### Gene Set Enrichment Analysis and Ingenuity Pathway Analysis

Gene set enrichment analysis was performed using GSEA v3.0, build 0160 (http://software.broadinstitute.org/gsea/) as described in (Subramanian et al., 2005). “Log2 ratio of classes” was used to rank changes in gene expression between developmental stages as well as between wild type and *DND_1_*^*Ter*/*Ter*^ samples. Normalized Enrichment Scores (NES) and FDR values were used to describe the magnitude of change and the significant enrichment of the gene sets respectively.

**Ingenuity Pathway Analysis.** Differentially expressed genes (DEGs) were assessed for statistical enrichment/depletion of specific pathways using the Qiagen Ingenuity Pathway Analysis (IPA) database (Application build: 470319M; Content version: 43605602). DEGs included those that were differentially expressed 1) between developmental stages within a *Ter* genotype class, and 2) between *Ter* genotype classes at a specific stage. The cutoff for inclusion in the IPA analysis was set at a differential expression FDR < 0.01.

**DND_1_ DO-RIP-seq.** DO-RIP-seq was performed as described in (Nicholson et al., 2017). Briefly, 5 - 15 cm^2^ plates of HEK_293_ cells were transfected with ~35 μg/plate FLAG-DND_1_ or untagged DND_1_ (as a negative control), grown for 24hrs, then harvested in PLB. Lysates were treated with 50 U/μg of MNase for 5 min at 30°C, and FLAG immunoprecipitations were performed. RNA from IPs was radiolabeled and run on a 10% TBE-Urea gel. RNA fragments from ~25-75 nucleotides were excised and extracted from the gel. cDNA libraries were prepared using the NEBNext Multiplexing Small RNA Library Prep Set for Illumina with 14-17 rounds of amplification, following the manufacturer,s protocol with the exception that the 5, RNA adapter was replaced with a custom 5, adapter compatible with UMI (Unique Molecular Identifier) tagging (Kivioja et al., 2011) (RNA adapter sequence GUUCAGAGUUCUACAGUCCGACGAUCNNNNN). In addition to DND_1_, negative control IPs input samples were also prepared for sequencing in a similar fashion using 17 rounds of amplification. An aliquot of the MNase-digested lysate prior to IP was set aside for generating input libraries. rRNA depletion was performed on input samples using the Epicenter Ribo-Zero Gold rRNA removal kit. Input RNA was size selected and libraries were prepared as described above for the IP samples. All experiments were performed as biological duplicates.

Libraries were sequenced using Illumina HiSeq 2500. Processing of data was performed as described in Nicholson et al. After removal of adapter and UMI sequences, reads were mapped to hg19 using TopHat2. Binding sites were identified using the binning and normalization procedure of Nicholson et al. (Nicholson et al., 2017), IPs were compared to input samples to identify sites enriched in DND_1_ IPs. Reads with a low pass filter of >3 reads and >1.5 fold enrichment over input were considered DND_1_ binding sites.

## Results

***DND_1_*^*Ter/Ter*^ mutant germ cells are delayed in the germ cell transcriptional profile relative to wild type at E_12.5_.** To investigate the changes in *DND_1_*^*Ter/Ter*^ mutant germ cells relative to wild type, 129T2-*DND_1_*^*Ter/+*^; *Oct4-EGFP* males and females were intercrossed, and ~600 germ cells were isolated by FACS from 4 independent homozygous and wild type fetal gonads at each of 3 stages, E_12.5_, E_13.5_ and E_14.5_, after germ cells have populated the gonads, and prior to teratoma detection (Fig. 1A; (Cook et al., 2011)). Sequencing was performed, reads were mapped to the genome, and a reads per kilobase transcript per million (RPKM) value was calculated for each transcript. RPKM values used in our analysis are provided with the capability to generate an expression graph for any gene, as a user-friendly resource for the community (Suppl. Table 1).

The *DND_1_* transcript was detected at similar levels in wild type and homozygous mutant germ cells at E_12.5_ (Fig. 1B), but showed a rapid decline at later stages (data not shown). The *Ter* mutation, predicted to lead to a premature stop codon (Youngren et al., 2005) is evident in the third exon of *DND_1_* and efficiently distinguishes wild type and mutant samples (Fig. 1B, blue arrow). The 129T2 variant of a SNP that differs between B6 and 129 was detected in the 3,UTR of all samples confirming the 129T2 strain background (red arrow).

*DND_1_* is expressed in both XX and XY germ cells from early migratory stages (Youngren et al., 2005). In *Dnd*^*Ter/Ter*^ mutants, many germ cells are lost prior to arrival in the gonad (Cook et al., 2009; Noguchi and Noguchi, 1985). Given that loss of RBPs in germ cells of other species results in transdifferentiation to somatic fate (Ciosk et al., 2006; Gross-Thebing et al., 2017; Updike et al., 2014), we investigated the possibility that *DND_1_*^*Ter/Ter*^ mutant germ cells were significantly altered by the time they populated the gonad, and did not retain their germ cell identity. The top two principle components account for 43% and 25% of the observed variation in the top 500 most variable protein-coding genes. Based on the variability captured by Principle Component (PC) 1, it appears that E_13.5_ mutant germ cells cluster more closely with E_12.5_ wild type germ cells while E_12.5_ mutant germ cells are delayed. Further, PC2 appears to stem mainly from expression variation in E_14.5_ *DND_1_*^*Ter/Ter*^ germ cells, suggesting that the transcription profile in these mutant germ cells continues to diverge from a normal trajectory (Fig. 1C). To investigate these differences further, we compared expression in *DND_1_*^*Ter/Ter*^ mutant germ cells with a list of genes specific to germ cells (Fig. 1D) and a list of genes that distinguish *male* germ cells (Suppl Fig. 2A) at E_12.5_ and E_13.5_. These lists include genes that characterize germ cells by their specific expression in this lineage, and those that characterize germ cells by their specific repression (eg. somatic genes)(Jameson et al., 2012). Based on heat map analysis, *DND_1_*^*Ter/Ter*^ mutant germ cells show an overall pattern similar to wild type male germ cells for both up- and down-regulated genes at E_12.5_ and E_13.5_. For example, both *Ddx4* (dead box helicase 4)(*Vasa*) and *Dazl* (deleted in azoospermia like) show a delay in activation, similar levels at E_13.5_, and a down regulation at E_14.5_, while *Stella (Dppa3* (developmental pluripotency associated protein 3)), a germ cell specific gene, and *Prdm1* (PR/SET domain 1) (*Blimp1*) show no significant differences from wild type at any stage (Fig. 1E). These data indicate that although *DND_1_*^*Ter/Ter*^ germ cells have a delayed gonadal differentiation program, they have a transcriptome similar to wild type germ cells soon after they arrive in the gonad.

**Figure 2.**
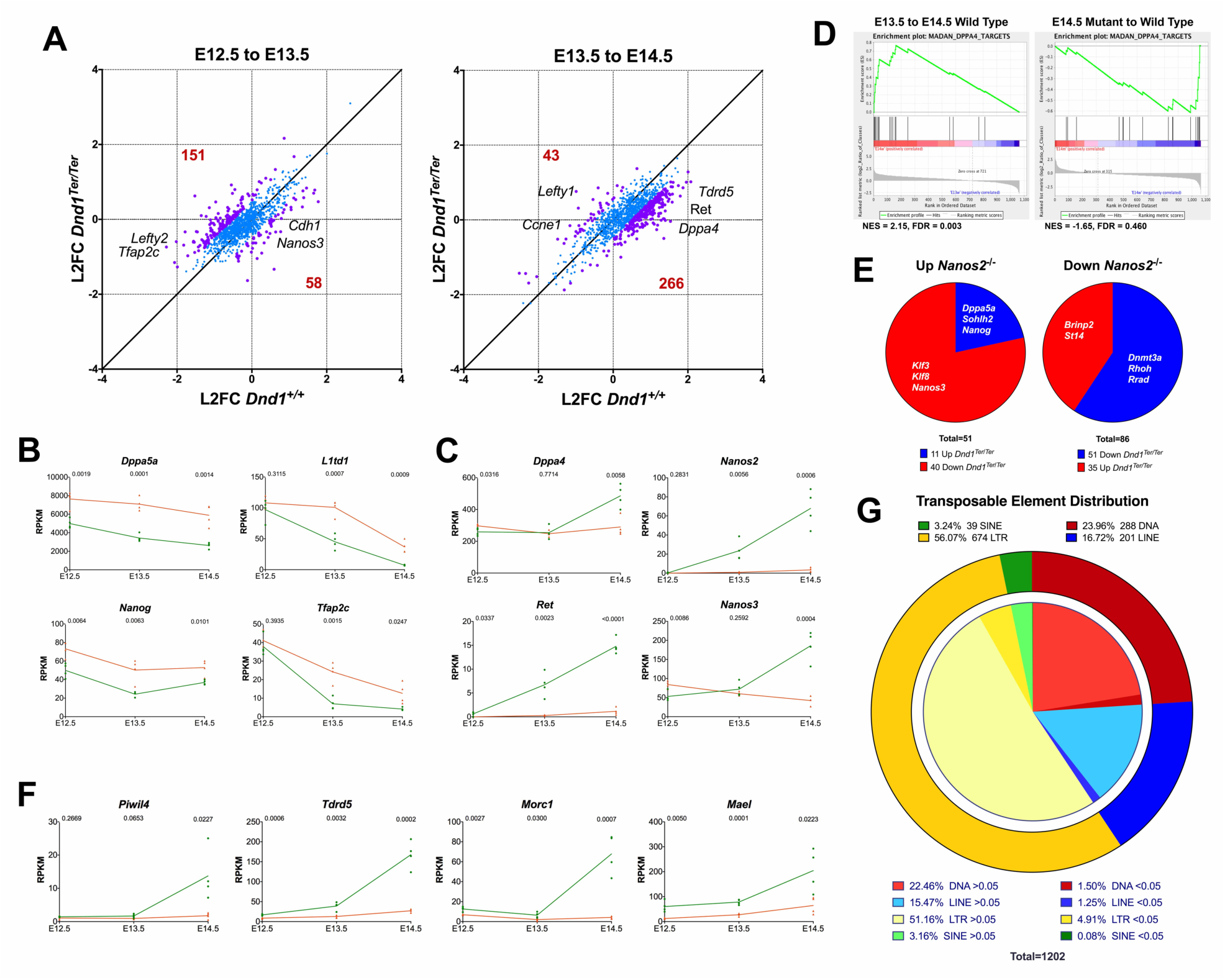
*DND_1_*^*Ter/Ter*^ mutant germ cells show delayed suppression of pluripotency, and a failure to activate the male differentiation program. **(A)** Scatter plots illustrating dynamic changes in expression of germ cell specific genes (data from Jameson et al., 2012) between mutant (Y axis) and wild type (X axis) during the transitions between E_12.5_-E_13.5_ and E_13.5_-E_14.5_. Between E_12.5_-E_13.5_ more genes (151) show a higher log_2_ fold change (L2FC) in mutants than in wild type, while only 58 genes lag behind. In the transition between E_13.5_-E_14.5_, the reverse is true: more genes lag behind wild type expression (266). Of the 43 genes that show a higher L2FC in this interval, many belong to cell cycle or elevated signaling pathways. Points with L2FC between mutant and wild type >0.5 in purple, <0.5 in blue. **(B)** Examples of 4 genes associated with pluripotency (*Dppa5a, L1td1, Nanog*, and *Tfap2c*) that show a delay in down-regulation in *DND_1_*^*Ter/Ter*^ mutant germ cells. **(C)** Examples of 4 genes associated with initiation of the male pathway (*Dppa4, NANOS_2_, Ret*, and *Nanos3*) that fail to be activated. **(D)** Consistent with this, based on GSEA analysis, targets of *Dppa4* are not enriched in mutant germ cells during the E_13.5_-E_14.5_ transition. **(E)** Some transcriptional changes in *NANOS_2_* mutants are discordant with changes in *DND_1_*^*Ter/Ter*^ mutants. Of genes up in *NANOS_2_* mutants, 11 behave the same (blue) and 40 (red) are down in *DND_1_*^*Ter/Ter*^. Of genes down in NANOS_2_ mutants, 51 are also down in *DND_1_*^*Ter/Ter*^ (blue) and 35 are up (red)(Saba et al., 2014). Only genes with significant L2FC are shown (p<0.05). See also Suppl. Table 3. **(F)** Examples of genes associated with suppression of transposable elements and the piRNA pathway activated in wild type but not mutants. **(G)** Although many silencers of repetitive elements were down-regulated, RepEnrich analysis showed that transposable elements did not exhibit significant aberrant activity at E_14.5_. Transposable element distribution organized by class: Outer ring: class distribution of elements represented in reads at E_14.5_ (yellow=LTRs; green=SINEs; Red=DNA transposons; Blue=LINEs). Inner circle divides specific classes into reads with significant differences between mutant and wildtype (bold colors: FDR<0.05; pale colors: FDR>0.05).

**Neither the female germ cell program nor other embryonic pathways are activated in XY *DND_1_*^*Ter/Ter*^ germ cells.** Because *DND_1_* expression becomes specific to the male pathway after germ cells enter the gonad (Youngren et al., 2005), we next investigated whether there was evidence for a shift to the female germ cell-specific program in *DND_1_*^*Ter/Ter*^ male germ cells by comparing expression in mutants to a list of genes specific to female germ cells (Jameson et al., 2012). A heat map comparison showed little evidence for ectopic activation of female pathway genes (Suppl. Fig. 2B). Based on a cumulative distribution plot, few female-specific genes were significantly altered in mutants at E_13.5_ (Suppl. Fig. 2C). In contrast to their pattern in wild type female germ cells, genes that are associated with entry into meiosis (eg. *Stra8* (stimulated by retinoic acid 8), *Sycp3* (synaptonemal complex protein 3)) were not upregulated in *DND_1_*^*Ter/Ter*^ mutant XY germ cells (Suppl. Fig. 2D). However, some genes that become female-specific by downregulation in male germ cells (eg. *Rhox9* (reproductive homeobox 9) and *Rhox6*) showed a less robust downregulation at the stage when male and female germ cell pathways diverge (E_12.5_) (Suppl. Fig. 2E). In general, these were genes associated with the pluripotent state of early germ cells.

Many genes associated with developmental pathways carry bivalent marks in germ cells (Lesch and Page, 2014). Because teratoma development is associated with the upregulation of developmental pathways, we investigated whether specific groups of bivalent genes showed upregulation in *DND_1_*^*Ter/Ter*^ germ cells. We found no evidence of lineage infidelity based on the activation of any subgroup of bivalent genes associated with embryonic pathways (Suppl. Fig. 3A), or significant differences in specific factors associated with differentiation (Suppl. Fig. 3B).

**Figure 3.**
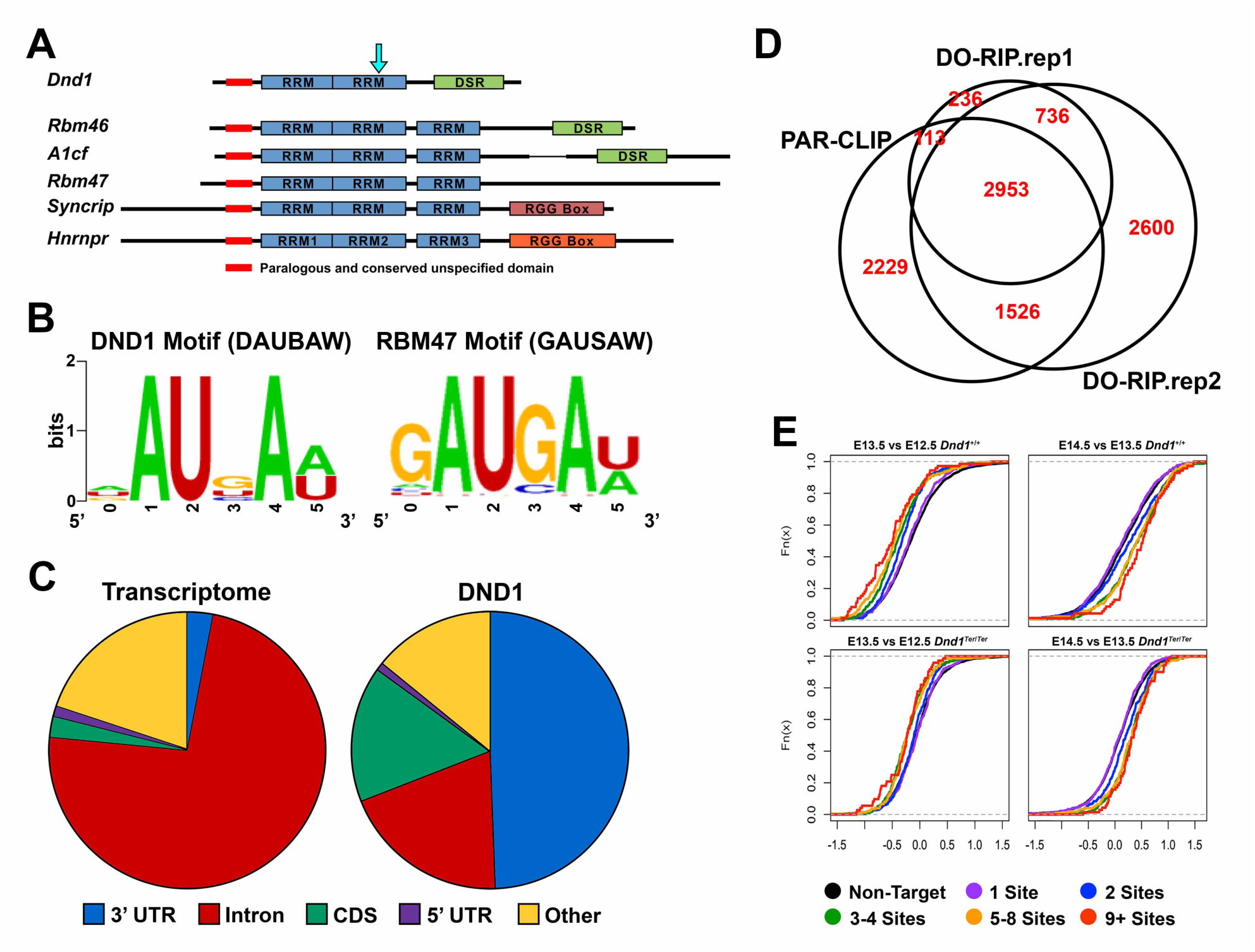
DND_1_ binds to the DATBAW motif preferentially localized to 3, UTRs, and regulates the up- and down-regulation of targets. **(A)** DND_1_ and five other RBPs (RBM46, A1CF, RBM47, SYNCRIP and HNRNPR) are part of a subfamily containing RNA recognition motifs (RRMs) sharing sequence similarity. Three of these subfamily member (DND_1_ included) also have similar C-terminal double stranded RNA binding domains (DSR) The black arrow indicates the position of the Ter mutation, partially truncating the second RRM and completely removing the DSR. **(B)** k-mer analysis was preformed to identify the most enriched motif in DND_1_ DO-RIP-Seq binding sites (http://weblogo.berkeley.edu), this motif is highly similar to a previously identified motif of a related RBP, Rbm47 (Ray et al., 2013). **(C)** The locations of DND_1_ binding sites were broken down into regions (3,UTR, intron, CDS, 5,UTR, and other) and compared to the proportion of the regions present in the expressed transcriptome. DND_1_ has a strong preference for binding 3,UTR sequences and a slightly weaker preference for coding sequence (CDS). **(D)** Comparison of DND_1_ binding sites identified in the two replicates of the present study and binding sites identified by PAR-CLIP in Yamaji et al., 2017 (Yamaji et al., 2017)**(E)** Cumulative distribution function (CDF) plots evaluating RNA abundance under four conditions (E_12.5_-E_13.5_, and E_13.5_-E_14.5_ in both mutant and wild type germ cells) with changes for transcripts classified by the number of DND_1_ binding sites (non-targets, 1 site, 2 sites, etc). In wild type germ cells RNA abundance of DND_1_ targets is more likely to decrease between E_12.5_ and E_13.5_ than for non-targets. In contrast, between 13.5 and 14.5 RNA abundance of DND_1_ targets generally increases. The effects are amplified by increasing the number of DND_1_ binding sites. In *DND_1_* mutant germ cells, RNA abundance changes for both E_12.5_-E_13.5_ and E_13.5_-E_14.5_ are similar for DND_1_ targets and non-targets.

### Homozygosity for the *DND_1_*^*Ter*^ mutation is associated with subtle changes in expression of many genes in male germ cells between E_12.5_-E_14.5_

Next we compared the effect of the *Ter* mutation on the temporal gene expression of germ cells during the transition between E_12.5_ and E_13.5_ and between E_13.5_ and E_14.5_. While most expressed genes retain a similar temporal pattern in the mutant compared to wild type over this period, hundreds are up- and down-regulated between E_12.5_-13.5 and/or E_13.5_-14.5 (Fig. 2A). More genes are up-regulated in mutant PGCs relative to wildtype during the first transition between E_12.5_-13.5 (151 up v. 58 down), whereas more genes are down-regulated in mutant PGCs during the second transition between E_13.5_-14.5 (43 up v. 266 down). Transcripts for many pluripotency genes are enriched in mutant PGCs throughout the time course (Fig. 2B). A large group of genes associated with the initiation of male germ cell development, including *NANOS_2_, Nanos3* (C2HC-type zinc finger 3), *Ret* (rearranged during transfection – a receptor tyrosine kinase), *Ddx25* (dead box helicase 25), and *Dppa4* (developmental pluripotency-associated protein 4), a marker of pluripotent cells required for differentiation (Madan et al., 2009) fail to be activated in mutants (Fig. 2C). Based on Gene Set Enrichment Analysis (GSEA), targets of DPPA4 (including *Gtsf1* (gametocyte specific factor 1), *Stk31* (serine threonine kinase 31), *Ddx4* (dead box helicase 4), *Iqcg* (IQ motif containing G), *Mael* (maelstrom spermatogenic transposon silencer), *Slc25A31* (solute carrier famly 25), *Rnf17* (ring finger protein 17), and *Mep1b* (meprin A subunit beta), Suppl. Table 2) are enriched in wildtype PGCs (Fig. 2D, left panel) but not in mutant PGCs (Fig. 2D, right panel) during the E_13.5_-E_14.5_ transition, further suggesting that the *Ter* mutation disrupts *Dppa4* expression and its downstream function in PGCs.

Given the critical roles of *NANOS_2_* in establishing the male pathway in mouse germ cells (Suzuki and Saga, 2008), we asked whether most transcriptional changes in *DND_1_*^*Ter/Ter*^ mutants could be accounted for by failure to activate *NANOS_2_* (Fig. 2C). To investigate this possibility, we compared the transcriptome changes between the two mutants. Although some genes changed in a similar manner (eg. genes associated with pluripotency), many genes exhibit opposing patterns (Fig. 2E). For example, *Ret* was not affected in *NANOS_2_* mutants whereas *Sycp3* and *Stra8* were elevated (Saba et al., 2014). In *NANOS_2_* mutants, *Nan*os3 was upregulated, whereas in *DND_1_* mutants, it was not (Fig. 2C; see Suppl. Table 3 and for full data set). This indicates that DND_1_ is upstream of both Nanos genes, and is consistent with the finding that *Nanos3* can partially rescue the *NANOS_2_* mutants (Saba et al, 2014).

### Transposable Elements and piRNA pathways were not strongly affected by E_14.5_, despite the failure to activate their repressors

Several genes involved in repression of transposable elements including *Piwil4* (piwi like RNA-mediated gene silencing 4), *Tdrd5* (tudor domain containing 5), *Morc1* (MORC family CW-type zinc finger 1), and *Mael* were either not activated or expressed at much lower levels in mutants (Fig. 2F). Proteins encoded by these genes normally silence LINE and other transposable elements. Our transcriptome data was based on oligo-dT priming, however some transposable elements contain polyadenylation sites (Lee et al., 2008). We interrogated the raw transcriptome data at E_14.5_ for activation of transposable/repetitive elements using RepEnrich, a program optimized to detect differential enrichment between two RNA-seq datasets (Criscione et al., 2014). Very few transposable element reads showed a significant difference between wild type and mutant germ cells (FDR < 0.05) (Fig. 2G). This trend of similarity between wild type and mutant germ cells was also observed when we interrogated repetitive elements and transposable elements more generally (Suppl. Fig. 4A,B).

**Figure 4.**
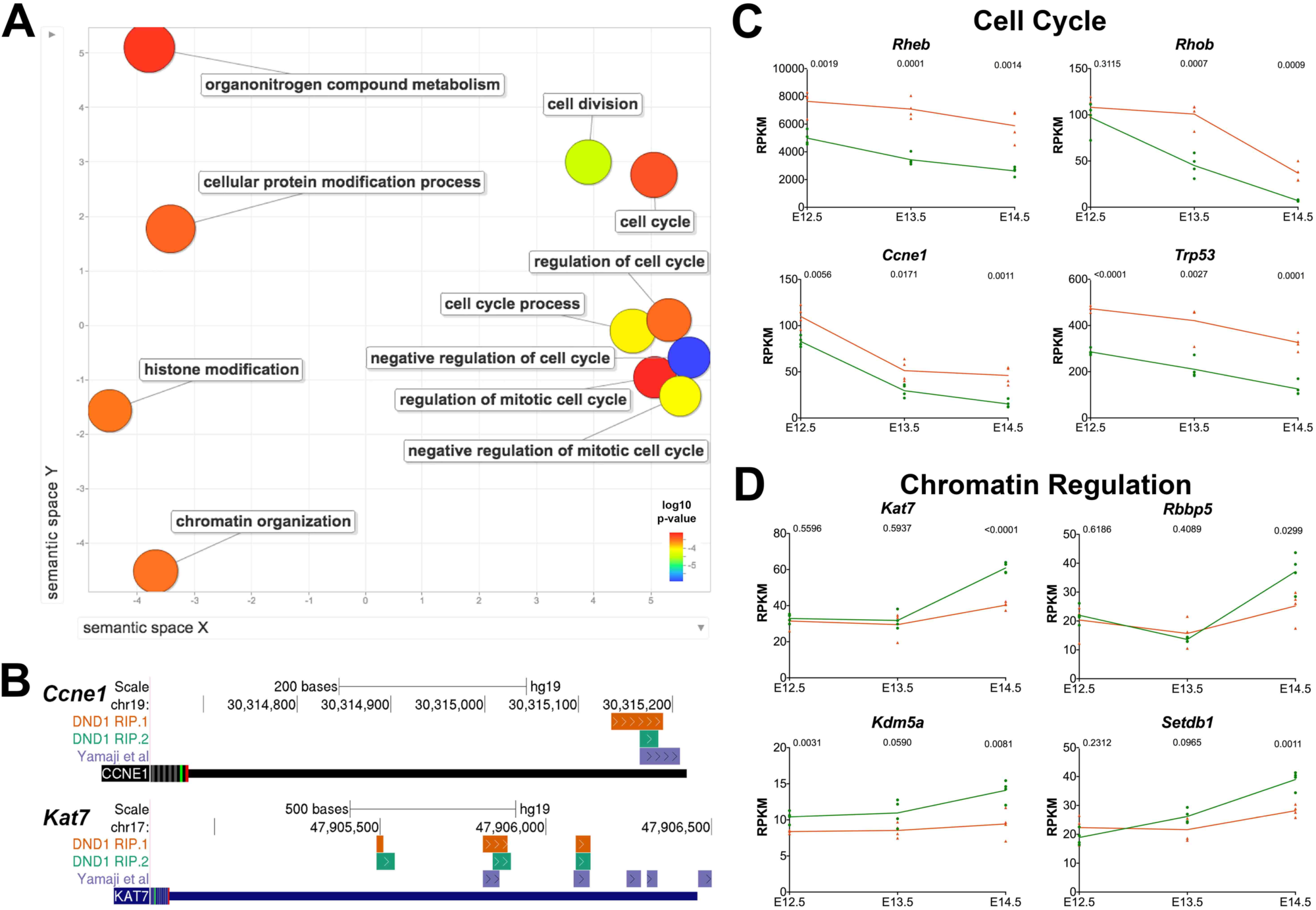
**DND_1_ targets include cell cycle genes and chromatin regulators, which are both up and down-regulated in mutant germ cells. (A)** Semantic similarity clustering of GO terms enriched in DND_1_ DO-RIP-seq targets indicated that the most common categories of targets of DND_1_ are genes associated with cell cycle and chromatin regulation. **(B)** Two examples of DND_1_ targets, *Ccne1* and *Kat7*. Binding sites for DND_1_ are present in the 3,UTR of *Ccne1* and *Kat7* in both DO-RIP-seq experiments and in the Yamaji dataset (Yamaji et al., 2017). **(C)** Examples of cell cycle gene targets that are up-regulated in mutants at E_14.5_ (*Rheb, Rhob, Ccne1*, and *Trp53*). **(D)** Examples of chromatin regulators that are down-regulated in mutants relative to wild type by E_14.5_ (*Kat7, Rbbp5, Kdm5a*, and *Setdb1*).

DND_1_ was previously reported to protect transcripts from miR470 (in zebrafish) and mir-1, miR-206, and miR-221 (in human cell lines) -mediated degradation (Kedde et al., 2007). Therefore, we predicted that targets of a mouse ortholog of these miRNAs such as miR302 (the ortholog of zebrafish miR470) would be down-regulated in the absence of DND_1_ protection. However, only *Lats2* (large tumor suppressor kinase 2) showed this behavior; other targets of miR302 were elevated in *DND_1_*^*Ter/Ter*^ germ cells relative to wildtype (Suppl. Fig. 5A). We used Ingenuity Pathway Analysis (IPA-Qiagen) to infer activation of individual miRNAs on a global basis. IPA compares the behavior of sets of miRNA targets among samples, calculates a p-value based on this overlap, and returns a Z-score reflecting the probability that a particular miRNA pathway is activated or repressed based on the expression of target genes. Based on IPA analysis, only one miRNA (mir223) cleared the activation cut-off of 2 between E13.4 and E_14.5_ in mutant cells (Suppl. Fig. 5B). However, investigation of the expression of predicted targets of mir223 revealed no consistent pattern (data not shown). Overall, we discovered no strong miRNA regulatory signature in this mRNA-seq analysis.

**Figure 5.**
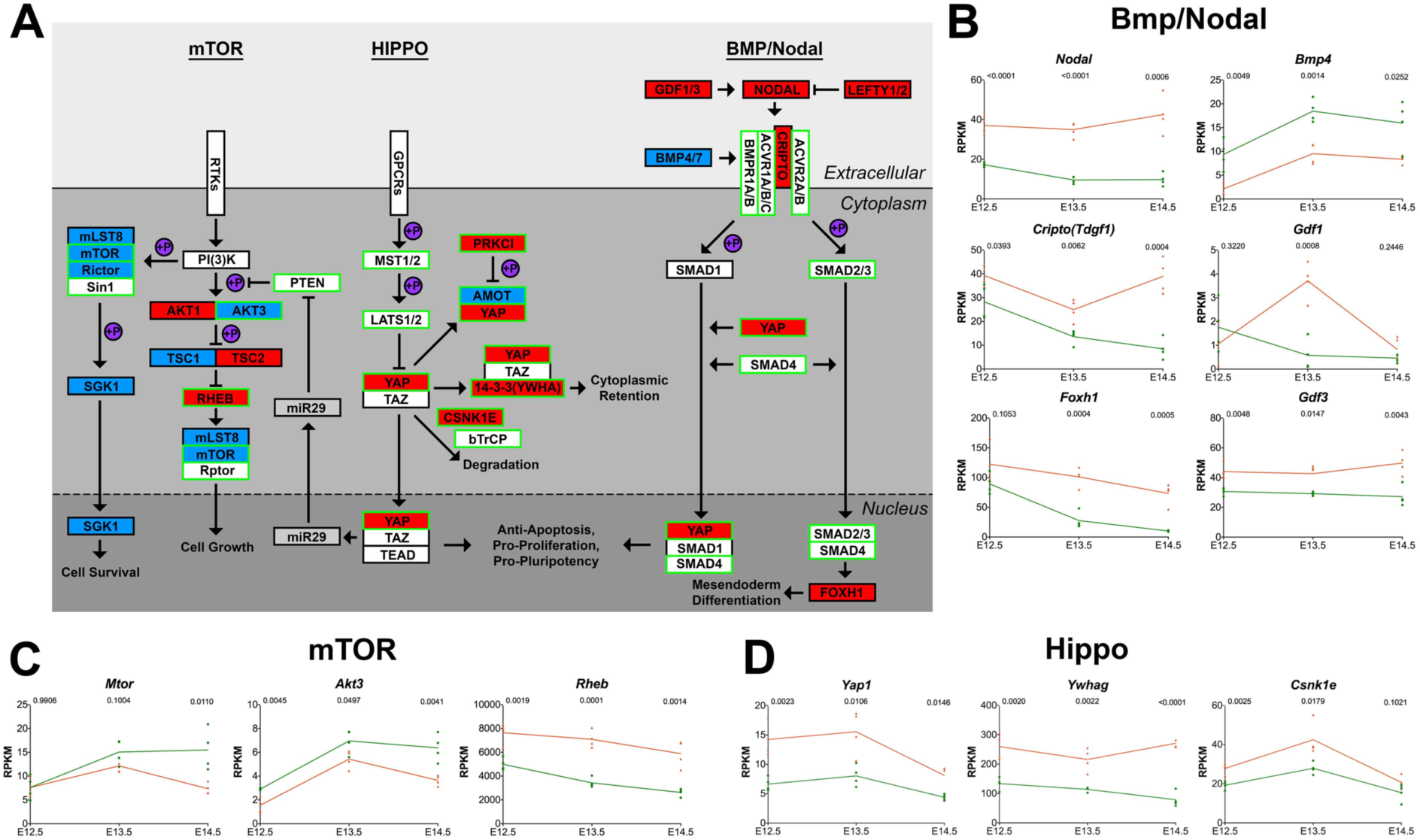
**In the presence of the *Ter* mutation in *DND_1_*, members of mTOR, Hippo and Bmp/Nodal pathways have altered expression with some transcripts indicated as direct targets of DND_1_. (A)** Interactions for mTOR, Hippo, and BMP/Nodal signaling pathways based primarily on KEGG analysis (Kanehisa et al., 2017) with input from primary literature (Beyer et al., 2013; Fuerer et al., 2014; Shimobayashi and Hall, 2014; Slagle et al., 2011; Spiller et al., 2017; Tanaka et al., 2007; Yu and Guan, 2013). Genes in pathways showing altered expression are colored blue for depression and red for elevation (RNAseq). Genes that map as transcript targets of DND_1_ based on RIP/ RIPseq are boxed in green. **(B,C,D)** Graphs for representative genes in pathways. **(B)** *Nodal, Cripto, Foxh1*, and *Gdf3* show strong up-regulation while *Bmp4* is down-regulated in the mutant transcriptome data. *Gdf1* has a spike in up-regulation after E_12.5_. **(C)** Genes mapping as DND_1_ targets in the mTor pathway including *Mtor* and *Akt3* are down-regulated while the target *Rheb* is strongly up-regulated in mutants. **(D)** Genes in the Hippo pathway, including *Yap1, Ywhag,* and *Csnk1e* are targets of DND_1_ that are up-regulated in mutants.

**Identification of DND_1_ direct targets.** We identified hundreds of genes that were up- and down-regulated between E_12.5_-14.5 in DND_1_ mutant PGCs (Fig. 2A). DND_1_ has been shown to both promote degradation of target transcripts (Suzuki et al., 2014; Yamaji et al., 2017) and to stabilize other target transcripts by competing for miRNA binding sites (Kedde et al., 2007; Zhu et al., 2011). We were interested in whether genes in both up- and down-regulated categories were direct targets of DND_1_.

*DND_1_* shares sequence similarity with genes encoding five other RNA binding proteins (*Rbm46* (RNA binding motif protein 46), *A1cf* (APOBEC complementation factor), *Rbm47, Syncrip* (synaptotagmin binding cytoplasmic RNA interacting protein) and *Hnrnpr* (heterogeneous nuclear ribonucleprotein R). All members of this subfamily contain three RNA recognition motifs (RRMs) except DND_1_ which contains two RRMs. Additionally, three members of the family, including DND_1_, contain a C-terminal double stranded RNA binding domain (DSR). (Fig. 3A). The *Ter* mutation generates a premature stop codon in the middle of RRM2 (arrow). To identify binding sites of DND_1_ we performed DO-RIP-seq (Nicholson et al., 2017) in HEK_293_ cells, using a tagged mouse DND_1_. The most enriched motif in DND_1_ binding sites was DAUBAW (D=A, G or U, B=C, G or U and W=A or U), a motif that is very similar to the known RNA motif of RBM47, its closest relative (Fig. 3B). The RBM47 motif defined by RNA-Compete (Ray et al., 2013) is GAUSAW (S=G or C and W=A or U), both the DND_1_ and RBM47 motifs contain AU in the second and third position and AW in the fifth and sixth, but differ in the first and fourth position where the recognition sequence is more degenerate. Although *Rbm47* expression declines sharply between E_12.5_ and E_14.5_, *Rbm46, Rbm47, Syncrip*, and *Hnrnpr* are all expressed in germ cells in the gonad, and could partially compensate for loss of DND_1_ (Suppl. Fig. 6).

Compared to their proportional representation across the whole transcriptome, DND_1_ binding sites were strongly over-represented in 3, untranslated regions (UTRs) and to a lesser extent in the coding sequence (CDS) (Fig. 3C). Targets identified by DO-RIP-seq largely agree with PAR-CLIP identified DND_1_ targets from Yamji et al, with 2953 mRNAs shared in common (Yamaji et al., 2017)(Fig. 3D). This represents a significant overlap (p<0.001) based on 1000 iterations of randomly shuffled binding sites matched for size and restricted to expressed 3,UTRs.

**Direct targets of DND_1_ are both up- and down-regulated in *DND_1_*^*Ter/Ter*^ mutants.** Because RIP assays were not conducted in germ cells due to the absence of appropriate antibodies or transgenic tags, we first identified the set of conserved targets expressed in germ cells (1347 out of 2953 targets). Next we investigated the relationship between number of DND_1_ binding sites in the target transcript (1 to 9+ binding sites) and its expression in wild type and 129T2 *DND_1_*^*Ter/Ter*^ mutant germ cells between E_12.5_-E_13.5_ and E_13.5_-E_14.5_. Cumulative distribution function (CDF) plots were generated comparing RNA abundance changes for transcripts classified by number of DND_1_ binding sites (non-targets, 1 site, 2 sites, etc). Changes in RNA abundance were evaluated in this manner for four conditions: E_12.5_-E_13.5_, and E_13.5_-E_14.5_ in both mutant and wild type germ cells. In wild type germ cells, the presence of more DND_1_ binding sites in a target transcript was correlated with a higher likelihood of down-regulation relative to transcripts with few or no target sites during the transition between E_12.5_-E_13.5_, while the opposite pattern was observed between E_13.5_-E_14.5_ – target transcripts with more DND_1_ binding sites were more likely to be up-regulated relative to non-targets. During this later transition, DND_1_ targets tend to change faster than the rest of the transcriptome (Fig. 3E). In 129T2-*DND_1_*^*Ter/Ter*^ mutants, DND_1_ targets behave more like non-targets, reflecting the effect of the *Ter* mutation on DND_1_ function.

Overall, among targets of DND_1_, cell cycle genes and chromatin regulators were over-represented (Fig. 4A; Table 4). Overlapping binding sites from all three RIP assays were present in the 3,UTR of the cell cycle gene, *Ccne1*, and throughout the CDS and 3,UTR of the chromatin regulator, *Kat7* (lysine acetyltransferase 7) (Fig. 4B). In addition to *Ccne1, Rheb* (Ras homolog enriched in brain) and *Rhob* (Ras homolog family member B), two GTP-binding proteins associated with active cell cycle were targets of DND_1_, and were elevated in the mutant transcriptome along with *Trp57* (*Cdkn1c*, cyclin dependent kinase inhibitor P57), which was not expressed in HEK_293_ cells (Fig. 4C). *Kat7* and other chromatin regulators, including *Setdb1* (set domain bifurcated 1), *Kdm5a* (lysine demethylase 5a), and *Rbbp5* (RB Binding Protein 5, Histone Lysine Methyltransferase Complex Subunit) mapped as targets, and all lagged behind wild type levels (Fig. 4D), as did many other chromatin regulators not expressed in HEK_293_ cells (Suppl. Fig. 7).

Many genes associated with signaling pathways were identified as targets of DND_1_, and were significantly changed in mutants (Fig. 5A). A large group of genes associated with the BMP signaling pathway, were elevated in mutant germ cells (Fig. 5B), as also reported by Yamaji (Yamaji et al., 2017). We identified elements of other signaling pathways elevated in mutants that map as targets of DND_1_, including *Ywhah* (tyrosine 3-monooxygenase/tryptophan 5-monooxygenase activation protein eta), *Ywhag* (tyrosine 3-monooxygenase/tryptophan 5-monooxygenase activation protein gamma), *Csnk1e* (casein kinase 1 epsilon), *Yap1* (yes-associated protein 1), and *Prkci* (protein kinase C lota) in the Hippo pathway (Fig. 5C). In contrast, transcripts for two central genes in the mTOR pathway that mapped as direct targets of DND_1_, *Rictor* (rapamycin-insensitive companion of mTOR) and *Mtor* (mammalian target of rapamycin), were down-regulated in mutants (Fig. 5D).

**Ingenuity Pathway Analysis identified apoptotic and cell cycle pathways as strongly affected in mutants.** We used Ingenuity Pathway Analysis (Qiagen) to identify curated pathways most strongly affected in the mutant transcriptome. This analysis resulted in the identification of Cellular Growth and Proliferation (482 genes) and Cell Death and Survival (433 genes) as the most affected pathways (Suppl. Fig. 8A-C). These results are consistent with previous findings that many *DND_1_*^*Ter/Ter*^ germ cells do not enter cell cycle arrest, but are eliminated through apoptotic pathways (Cook et al., 2009; Dawson et al., 2018; Yamaji et al., 2017).

## Discussion

Germ cells are an important model to investigate how stem cells embark on a highly specific differentiation pathway while restraining their potential for spontaneous differentiation or tumorigenesis. While most germ cells appear to strictly control this potential and successfully navigate this transition, a small subset of 129T2 *DND_1_*^*Ter/Ter*^ mutant germ cells fail this test and spontaneously differentiate into teratomas - and in so doing provide a tool to investigate the genetic regulation of germ cell differentiation. What are the critical roles of DND_1_, and why does mutation in this RBP lead specifically to teratoma development in a subset of mutant germ cells? To address these questions, we investigated the binding targets of the DND_1_ protein and analyzed the transcriptome of *DND_1_*^*Ter/Ter*^ germ cells relative to wild type over the critical period following residency in the gonad but prior to overt transformation. We found that DND_1_ down-regulates genes associated with pluripotency, including multiple components of the Hippo and Bmp pathways, mediates levels of many genes involved in the control of cell cycle, and is critical to initiate male germ cell differentiation, which involves the activation of many chromatin regulators.

DND_1_ is expressed from the time germ cells are allocated at the base of the allantois, and as a result of its early role in germ cell development, many germ cells undergo apoptosis, or are otherwise lost, prior to arrival in the gonad (Cook et al., 2009; Noguchi and Noguchi, 1985). However, this transcriptome analysis reveals that overall, the transcriptome of the group of mutant germ cells that survive to populate the gonad at E_12.5_ is similar to wild type male germ cells, albeit somewhat delayed.

DND_1_ likely performs several roles in posttranscriptional regulation. E_13.5_ is a clear transition point between the embryonic and fetal programs: functions of DND_1_ may segregate into early exit from the pluripotency program, involving the degradation of transcripts for many pluripotency genes, and subsequent activation of transcripts associated with the male pro-spermatogonia pathway. Although we could not detect pluripotency targets in HEK_293_ cells, Matin and co-workers previously reported *Oct4, Nanog, Sox2, LIN28* (LIN28 family RNA-binding protein), *Bax* (BCL2 Associated X apoptosis regulator), and *Bclx* (B-cell lymphoma-estra large, apoptosis regulator) as targets of DND_1_ in ESCs (Zhu et al., 2011). A large group of chromatin regulators mapped as targets of DND_1_ in HEK_293_ cells, and all of these genes were strongly down-regulated in *DND_1_*^*Ter/Ter*^ mutants. However, it is unclear whether this down-regulation is the result of post-transcriptional control or a failure to launch the male-specific differentiation program.

There is strong evidence that DND_1_ interacts with the CNOT degradation complex, and provides target specificity for NANOS_2_ (Suzuki et al., 2014; Yamaji et al., 2017). Because *NANOS_2_* is not expressed in mutants, we asked whether the *DND_1_*^*Ter/Ter*^ mutant transcriptome phenocopied the *NANOS_2_* mutant transcriptome (Saba et al., 2014). We found that many pluripotency genes showed a similar delay in down regulation during the early transition (E_12.5_-E_13.5_). However, *NANOS_2_* mutant germ cells enter meiosis as though they have lost their male identity (Suzuki and Saga, 2008), whereas *DND_1_*^*Ter/Ter*^ germ cells do not. Key differences may be the up-regulation and partial compensation by *Nanos3* in *NANOS_2_* mutants, and/or the retention of *Ret*. Neither *Nanos3* nor *Ret* is expressed in *DND_1_*^*Ter/Ter*^ germ cells.

A key finding of this study is the global identification of many chromatin regulators that are activated in wild type male germ cells between E_13.5_-E_14.5_, and the observation that nearly all of these fail to be activated in *DND_1_*^*Ter/Ter*^ mutants. If the role of these chromatin regulators is to stabilize the spermatogonial genome and silence somatic differentiation pathways, as has often been proposed (Dawson et al., 2018; Gu et al., 2018; Heaney et al., 2012), the failure to activate these genes could explain the tendency of mutant germ cells to initiate differentiation to multiple cell types characteristic of teratomas. The failure to activate chromatin regulators was not reported in the original *NANOS_2_* mutants (Saba et al., 2014). However, heterozygosity for a CRISPR/*Cas9-*mediated knock out of *NANOS_2_* on the 129/SvJ background led to a decrease in DNMT3L expression, a defect in re-methylation of the male germ cell genome, elevated levels of *Line-1*, and an increase in teratoma incidence (Dawson et al., 2018).

Many genes associated with the cell cycle are targets of DND_1_ and are dysregulated in mutants (this study;(Cook et al., 2011; Yamaji et al., 2017). Several signaling pathways, including Kit, Fgf, Nodal, and Hippo are also elevated and could be responsible for driving ongoing proliferation. The association of FOXH1 with the YAP/SMAD complex is believed to drive the switch from pluripotency to differentiation of mesendoderm (Slagle et al., 2011), but further studies would be required to determine whether this finding has relevance for the differentiation of cell types in teratomas. A failure to exit active cell cycle and enter G0 arrest is a characteristic of mutant germ cells and teratoma formation, but whether this failure is causal remains unclear.

Based on the failure to activate many repressors of transposable elements such as *Piwil4, Morc1, Mael*, and *Tdrd5*, we anticipated a strong activation of LINES and other transposons. However, at the latest stage of our analysis (E_14.5_), we did not detect significant changes in this category. Similarly, despite the absence of many epigenetic regulators that normally characterize the male germ cell transcriptome at these stages, we found no evidence of lineage infidelity. One possibility is that E_14.5_ (the latest stage of this analysis) is too early to detect these changes at the population level. There is significant variability in the response of individual germ cells to the *DND_1_*^*Ter*^ mutation. *DND_1_*^*Ter*^ mutant germ cells transform and form teratomas at a much higher rate than wildtype germ cells. However, even though the “per mouse” teratoma rate is high, the “per germ cell” transformation rate is still quite low. Most *DND_1_*^*Ter/Ter*^ germ cells undergo apoptosis, as evidenced by the strong footprint of cell death and cell cycle disruption at the population level. A molecular explanation for this cell heterogeneity remains elusive. For stem cell biology, it would be very instructive to know the steps involved in the transition from a germ cell to a teratoma. Dawson and colleagues have recently suggested that this involves a transition through a primed pluripotent EC cell state (Dawson, 2018). The fact that only a few germ cells form teratomas in a *DND_1_*^*Ter/Ter*^ testis, while other germ cells nearby either die or continue to express normal germ cell markers (Cook et al., 2011; Dawson et al., 2018), suggests that the trigger must be a threshold event that is influenced by cell autonomous fluctuations of factors in germ cells and/or by the microenvironment in the testis of some strains. Teratoma formation likely competes with apoptotic pathways that are strongly activated at all stages of our analysis. In this study, we have identified many targets of DND_1_ and captured their transcriptional changes in *DND_1_*^*Ter/Ter*^ germ cells. However, the specific transcriptional events in the rare cells undergoing the transition to teratoma may be obscured by the transcriptomes of the large number of cells that are not. We plan to use a single cell analysis in future experiments to resolve this problem.

## Acknowledgments

We would like to thank the teams in the FACS and Genomics Cores at Duke, especially Mike Cook, Nicolas Devos, and Olivier Fedrigo, who helped with these experiments and their analysis. We are also grateful to Alex Bortvin, who provided information for the RepEnrich analysis, to Joe Nadeau, who supplied the mice that founded our *DND_1_*^*Ter*^ colony, and to past and present members of the Capel lab, especially Jordan Batchvarov, who contributed to the management of the strain over the last 10 years and Mike Czerwinski, who transferred bioinformatics expertise to JG. This work was funded by a grant to BC from NIH (GM087500), Bridge Funding from DUMC, and a grant to JK from NCI (CA157268) and Bridge Funding from DUMC.

## Supplemental Figure Legends

**Suppl. Table 1.** RNAseq gene expression graphing. We provide an excel file with **(A)** the RPKM values used in our analysis and **(B)** the graphic format used to display gene expression in wild type and *DND_1_*^*Ter/Ter*^ mutants over time. The user can copy any from **(A)** with values into row 2 of **(B)** to generate the graph for that gene.

**Suppl. Table 2.** The list of changed genes defined by GSEA as part of the “DPPA4 Targets”.

**Suppl. Table 3.** List of genes that are **(A)** up-regulated or **(B)** down-regulated in *NANOS_2_* mutants, annotated for activity in *DND_1_*^*Ter/Ter*^ mutants.

**Suppl. Table 4. (A)** List of all genes (expressed or not expressed in HEK_293_ cells), indicating genes identified in DO-RIP and/or PAR-CLIP (Yamaji et al., 2017). **(B)** List of all binding sites enriched in DO-RIP-seq. **(C)** List of genes with reproducible 3,UTR binding sites enriched in both DO-RIP-seq experiments. This table was used to label targets in Figure 5A.

---

**Suppl. Fig. 1.**
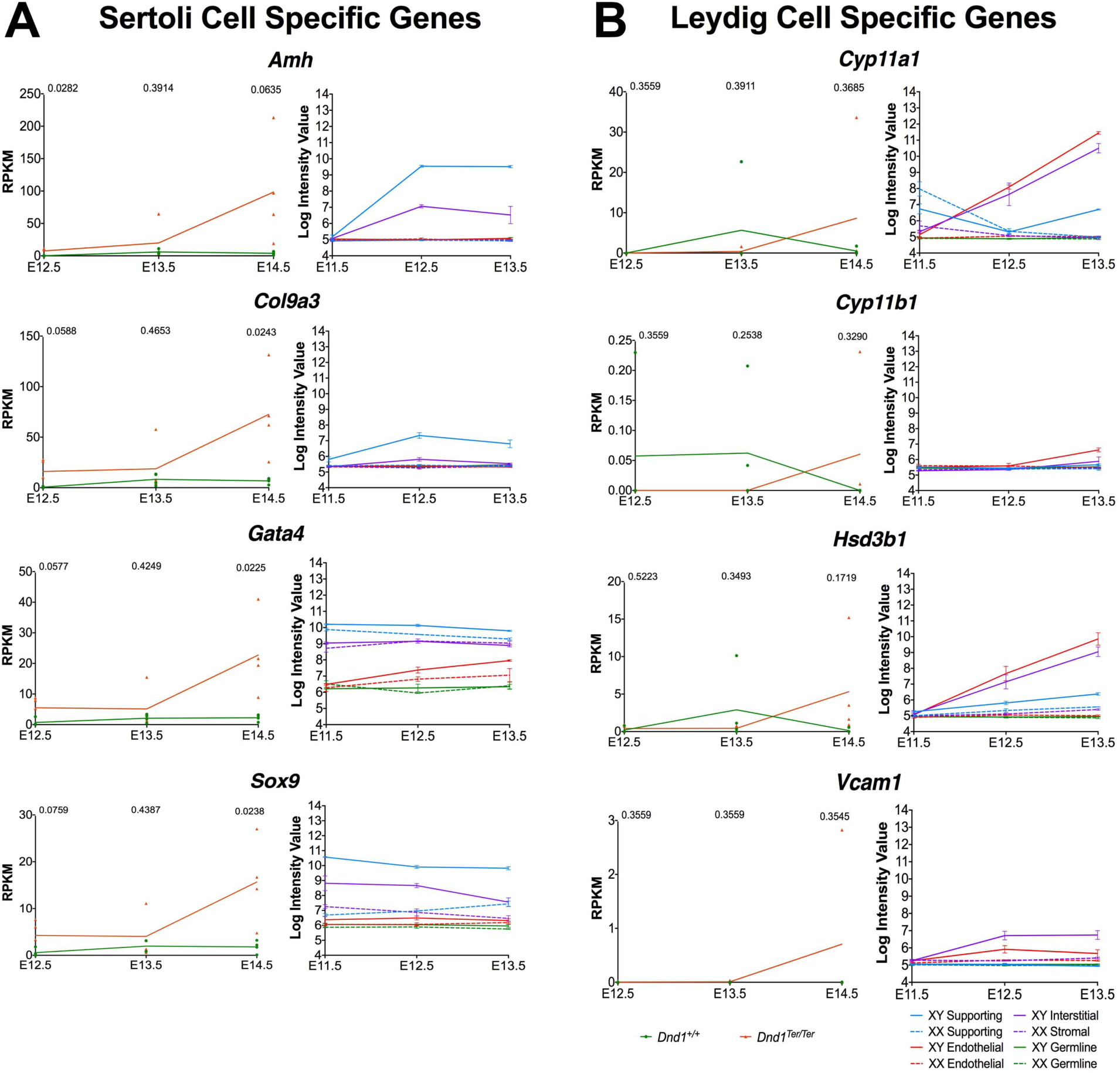
Examples of genes detected in the *DND_1_*^*Ter/Ter*^ transcriptome data that are specific to Sertoli or Leydig cell lineages in the gonad based on the Jameson data. (Jameson et al., 2012). Typically, expression of these genes spikes in the E_14.5_ *DND_1_*^*Ter/Ter*^ data, a stage when *DND_1_*^*Ter/Ter*^ germ cell numbers are reduced and expression of Oct4-GFP declines, which may have led to errors in threshold settings during FACS. Graphs of transcriptome data from each cell type in the gonad performed at E11.5, E_12.5_ and E_13.5_ are shown for comparison. The germ cell lineage is green; supporting cells are blue; interstitial/stromal cells are purple; endothelial cells are red.

**Suppl. Fig. 2.**
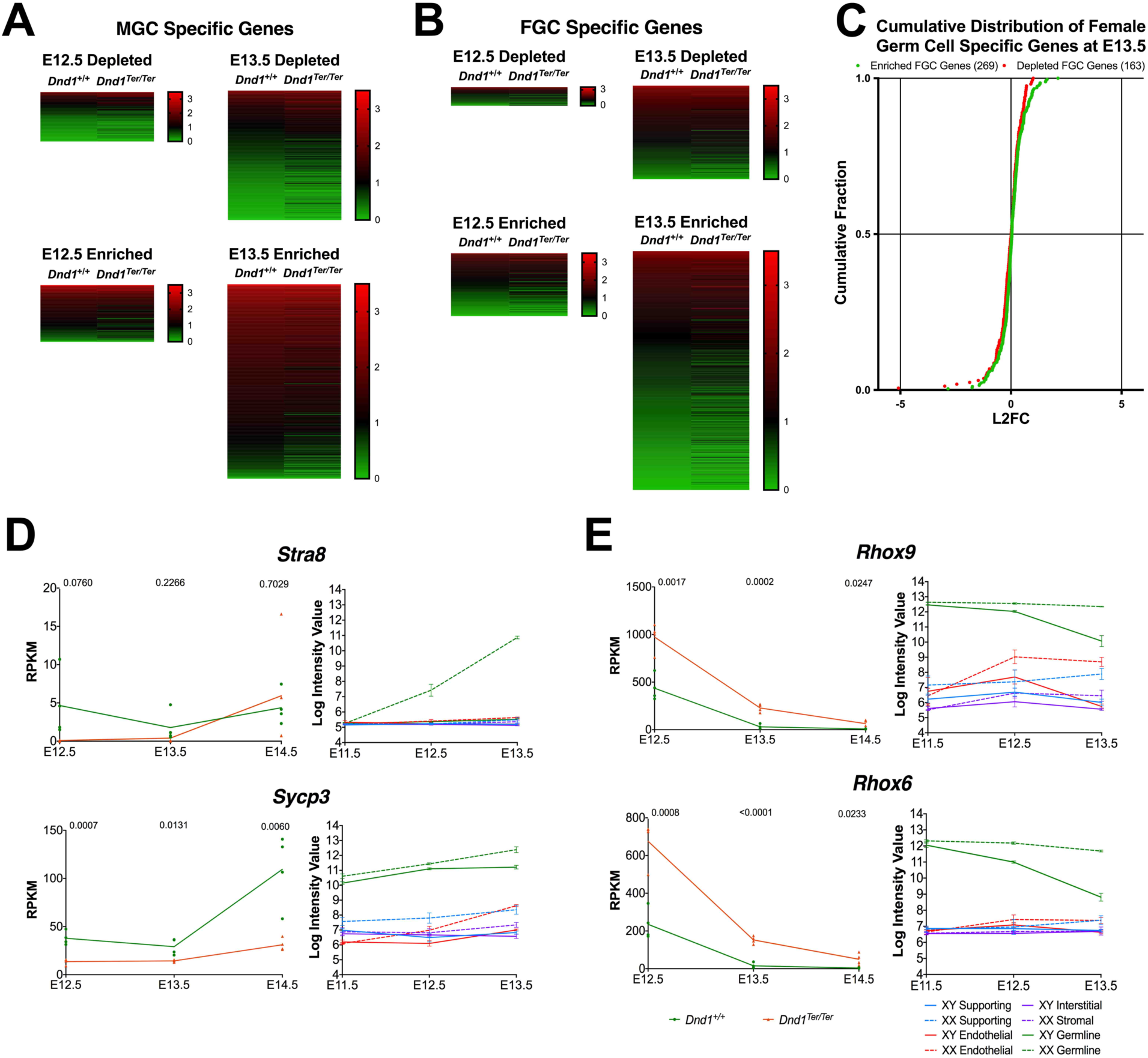
Male germ cells do not acquire a female (or meiotic) identity, but retain expression of pluripotency genes shared with female germ cells. **(A)** Heat map analysis showing that *DND_1_*^*Ter/Ter*^ male germ cells retain a profile very similar to wild type at E_12.5_ and E_13.5_ with respect to genes that are specifically depleted in male germ cells, and genes that are specifically enriched. **(B)** Similarly, heat maps comparing expression of female-specific genes in XY *DND_1_*^+/+^ to XY *DND_1_*^*Ter/Ter*^ samples show few changes for depleted or enriched genes at E_12.5_ or E_13.5_. **(C)** A cumulative distribution plot of the log2 fold change between mutant male germ cells and wild type male germ cells at E_13.5_ for female germ cell enriched (green) or depleted (red) genes. Genes typically enriched (green) or depleted (red) in female germ cells show no clear bias for the direction of change in mutant germ cells. **(D)** Genes specifically upregulated as female germ cells enter meiosis (*Stra8* and *Sycp3*) were not activated by E_14.5_ in *DND_1_*^*Ter/Ter*^ mutants. Graphs to the left show expression of the gene in all gonadal lineages (green broken line = XX germ cells; green solid line = XY germ cells; data from Jameson et al., 2012). **(E)** *Rhox9* and *Rhox6*, which normally become specific to XX germ cells by abrupt down-regulation in male germ cells, show a delay in down-regulation similar to other pluripotency genes.

**Suppl. Fig. 3.**
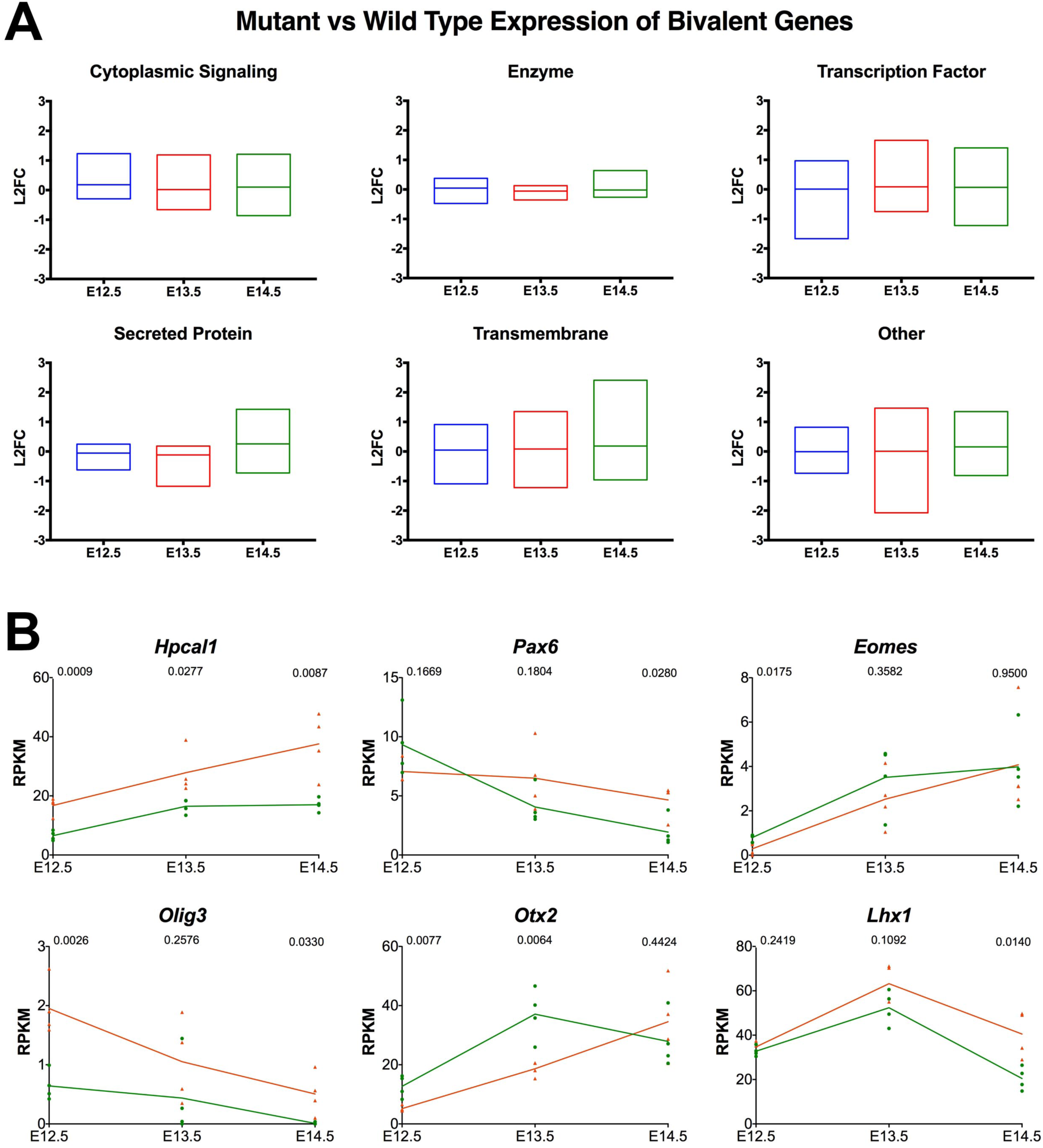
*DND_1_*^*Ter/Ter*^ germ cells show no evidence of lineage infidelity between E_12.5_-E_14.5_. Despite the failure to activate many chromatin regulators in *DND_1_*^*Ter/Ter*^ mutants, **(A)** expression of bivalent genes grouped by category (cytoplasmic signaling proteins, enzymes, transcription factors, secreted proteins, transmembrane proteins, and other) was not elevated, and **(B)** with the exception of *Hpcal1,* neither neuronal genes, nor other transcription factors associated with somatic cell differentiation (*Pax6, Eomes, Olig3, Otx2*, and *Lhx1*) were significantly elevated in mutants.

**Suppl. Fig. 4.**
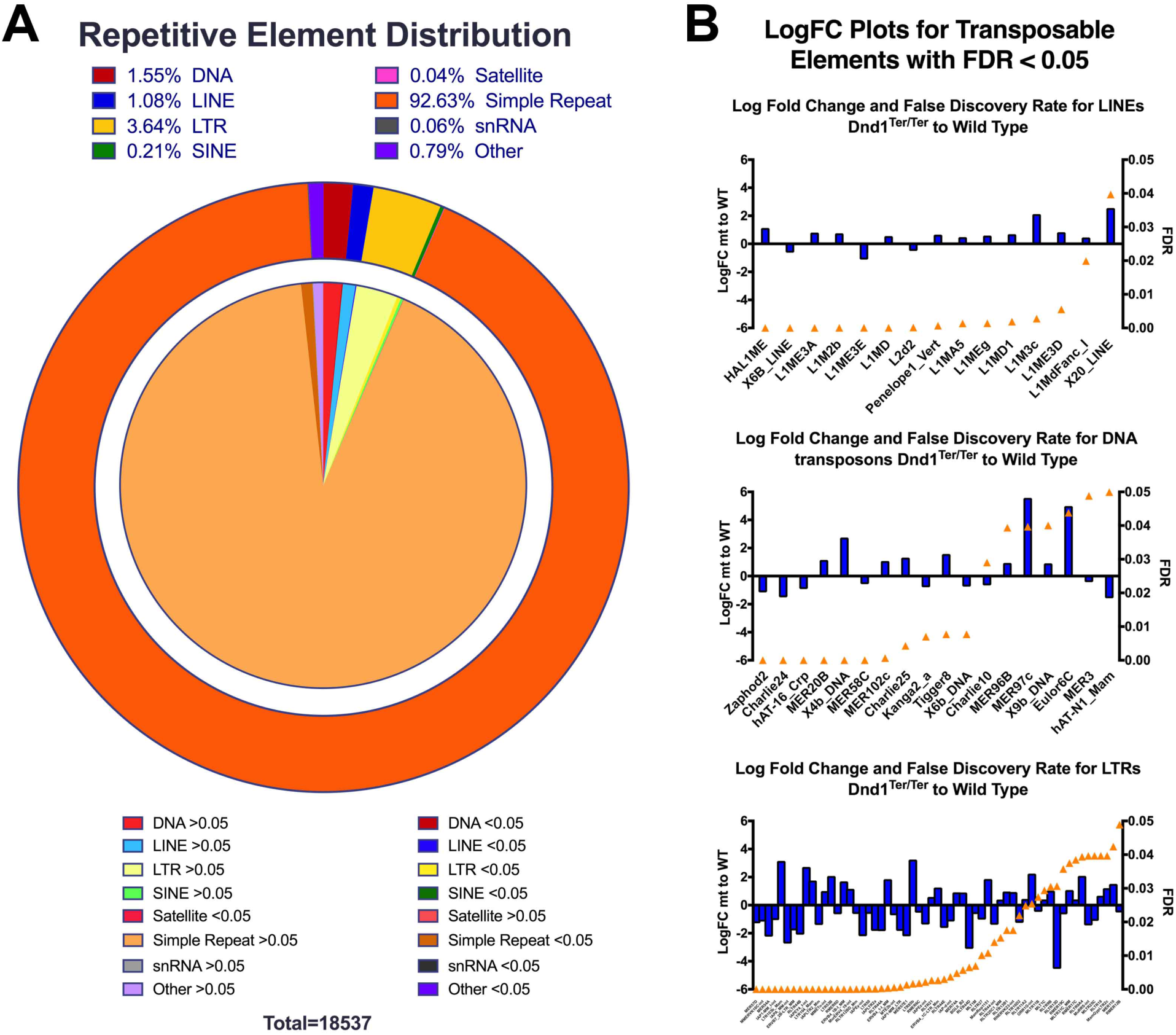
Repetitive elements in general do not exhibit aberrant activity. A survey of repetitive element distribution organized by class. Outer ring: class distribution of elements represented in reads at E_14.5_ (Red=DNA transposons; Blue=LINEs; gold=LTRs; green=SINEs; Magenta=satellite; Orange=simple repeats; Dark Green=snRNAs; Purple=other). Inner circle divides specific classes into reads with significant differences between mutant and wildtype and reads without significant differences between mutant and wildtype (bold colors: FDR<0.05, significant; pale colors: FDR>0.05). **(B)** Waterfall plots of Log Fold Change (LogFC) between mutant and wildtype in each transposable element class (LINEs, LTRs, and DNA transposons) with FDR<0.05 (significant). LogFC shown to left of plots; FDR shown by orange triangle placement relative to right scale on each graph.

**Suppl. Fig. 5.**
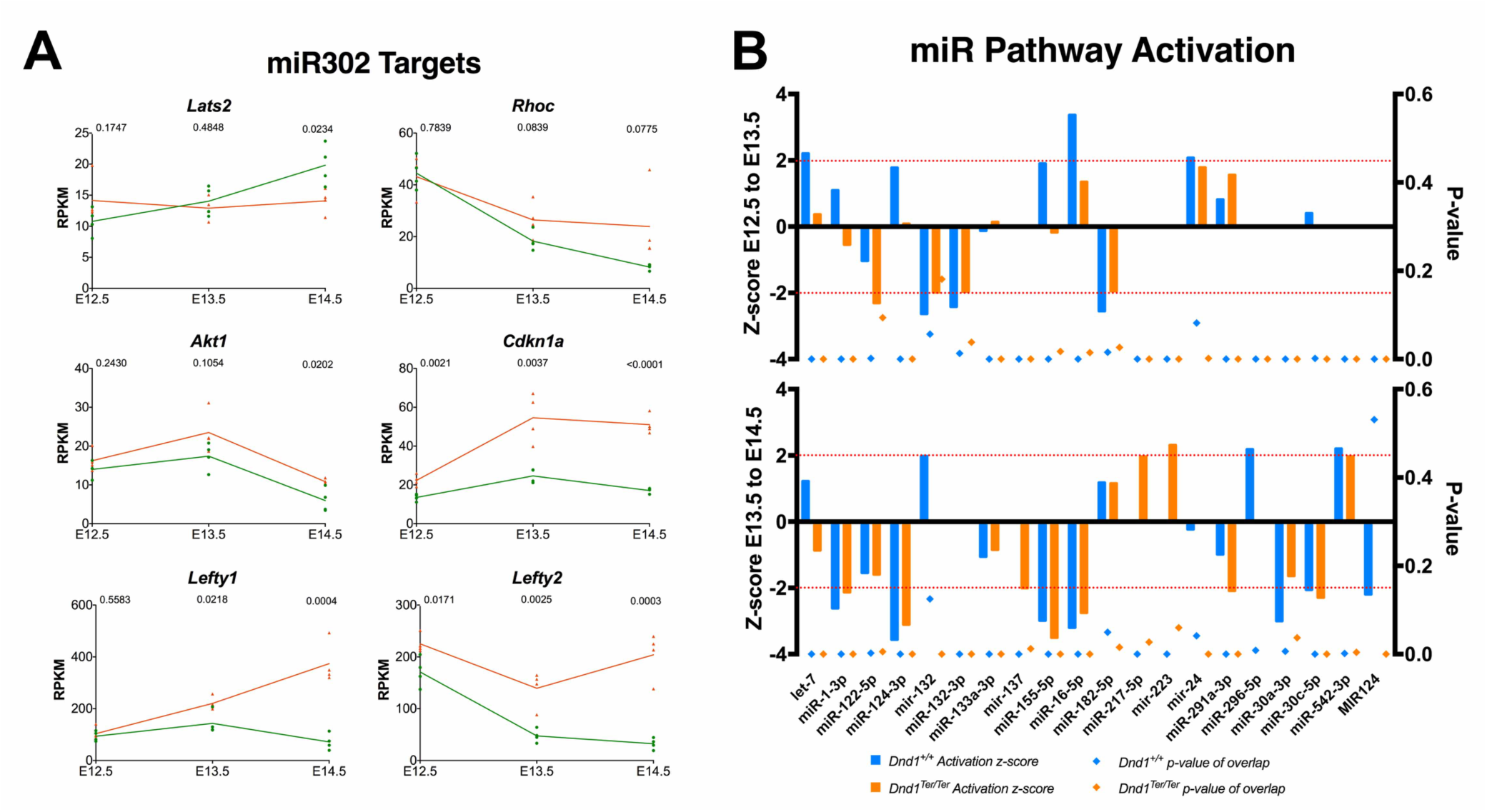
In general, miRNAs did not show activation based on downstream target analysis. **(A)** Although DND_1_ is believed to block degradation of zebrafish miR430 targets, with the exception of *Lats2*, targets of the orthologous miRNA in mice (miR320) are not down-regulated in mutants as predicted. **(B)** Waterfall graphs of Ingenuity Pathway Analysis (Qiagen) of miRNA activation based on expression changes in predicted targets at E_12.5_-E_13.5_ (top) and E_13.5_-E_14.5_ (bottom) for wild type (blue) and mutant (orange) samples. Only miRNA pathways with an activation Z-score of >2 or <-2 (left y-axis) in at least one transition period for mutant or wild type data are shown. P-values are plotted on the right Y-axis and represented as diamonds.

**Suppl. Fig. 6.**
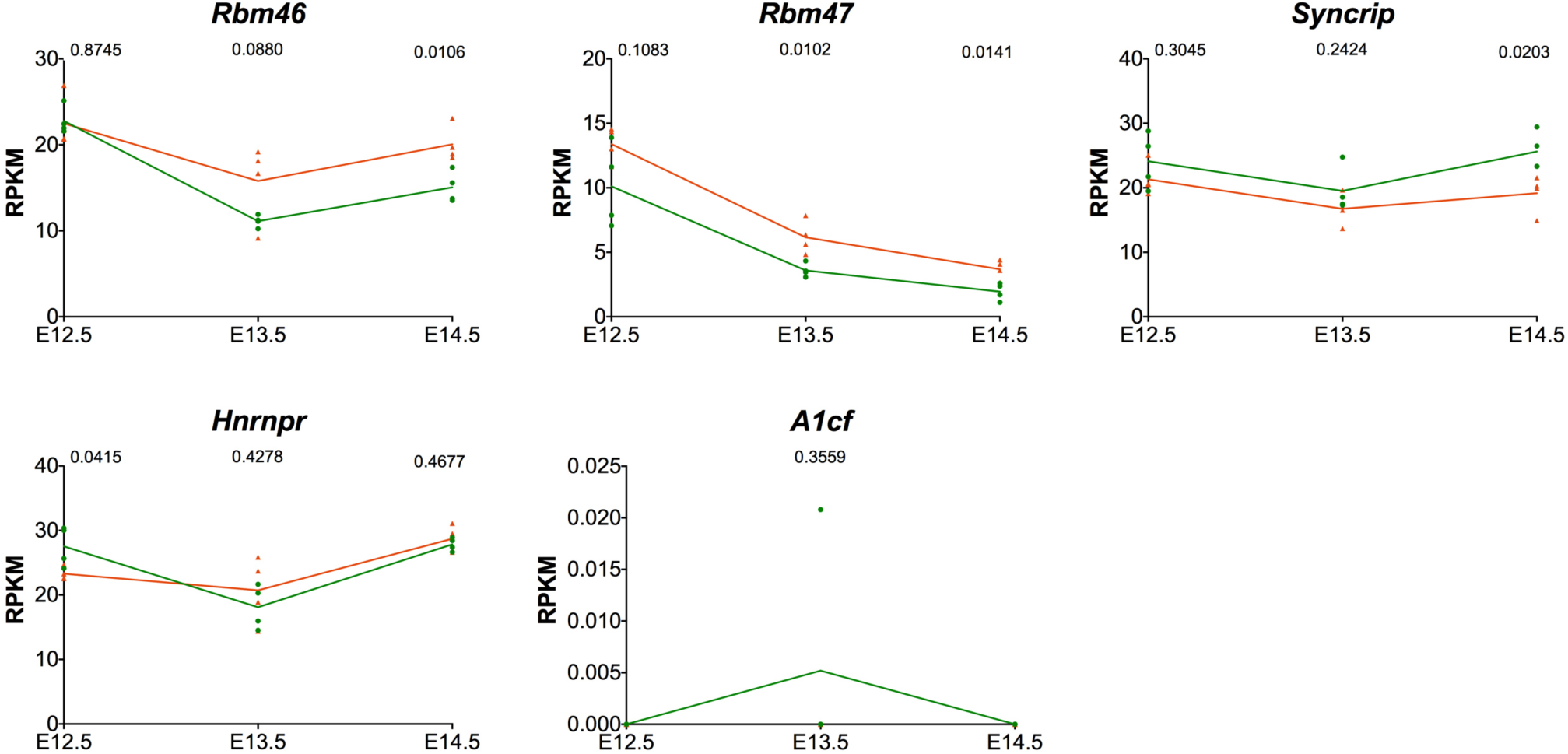
Four RBPS closely related to DND_1_ are expressed in germ cells. Four of the five RNA binding proteins (*Rbm47, Rbm46, Syncrip, Hnrnpr*, but not *A1cf*) related to DND_1_ (Fig. 3) are expressed in male germ cells during the transcriptome timecourse.

**Suppl. Fig. 7.**
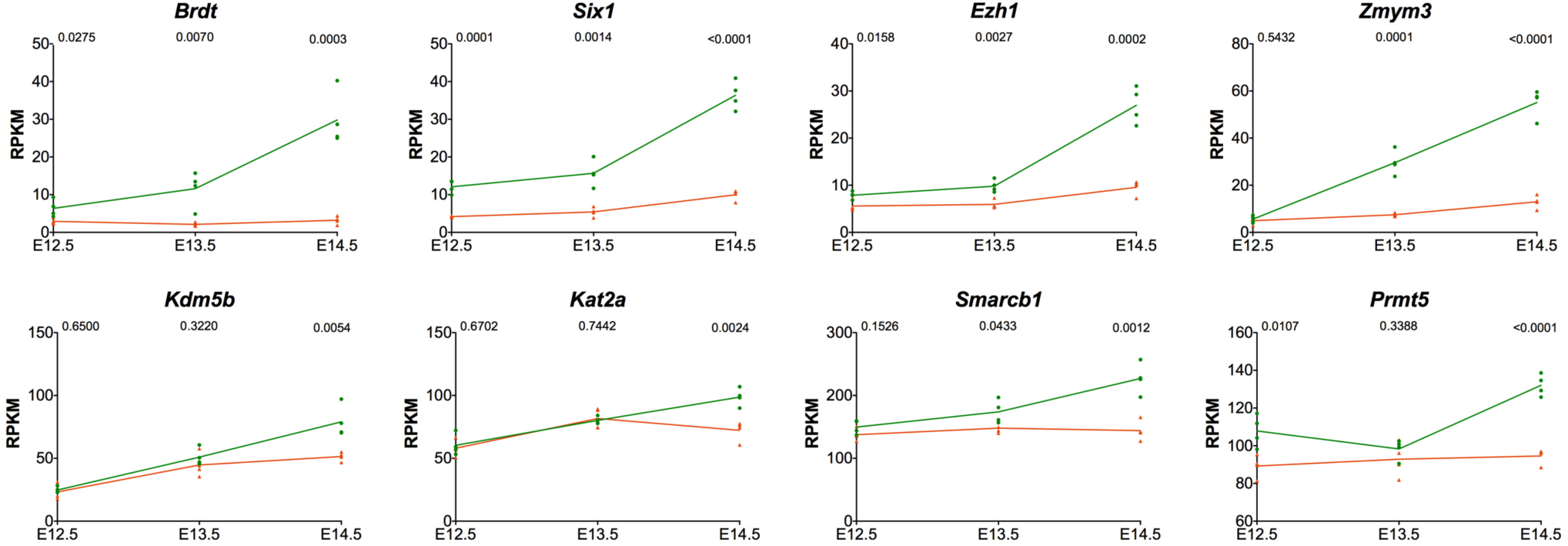
Many chromatin regulators that are not expressed in HEK_293_ cells, thus did not map as targets of DND_1_, fail to be activated in mutants. Nine additional chromatin regulators (*Brdt, Six1, Ezh1, Zmym3, Kdm5b, Kat2a, Smarcb1*, and *Prmt5*) are all expressed at lower levels in *DND_1_*^*Ter/Ter*^ mutants likely reflecting a failure to activate the male pathway.

**Suppl. Fig. 8.**
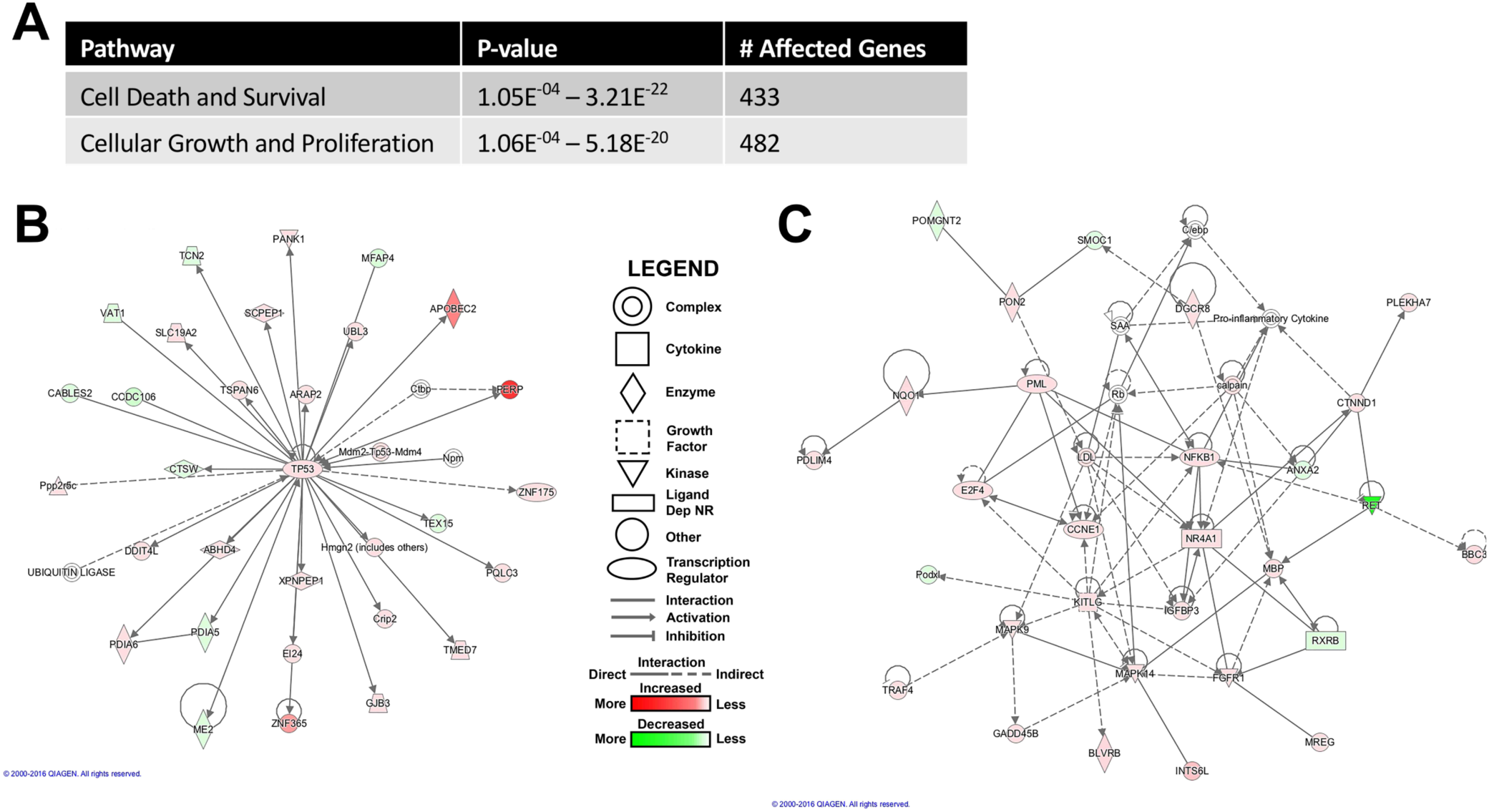
Cell death and cell cycle pathways are strongly affected in *DND_1_*^*Ter/Ter*^ mutants. **(A)** Ingenuity pathway analysis identified cell death (433 genes) and cell cycle (482 genes) pathways as the most strongly affected in *DND_1_*^*Ter/Ter*^ mutants. **(B)** Network diagram centered on *Tp53*, depicting the relationship of many cell death and survival genes. **(C)** Network diagram centered on *Ccne1*, depicting the relationship of many cell cycle genes, most of which are up-regulated.

**Suppl. Fig. 9.**
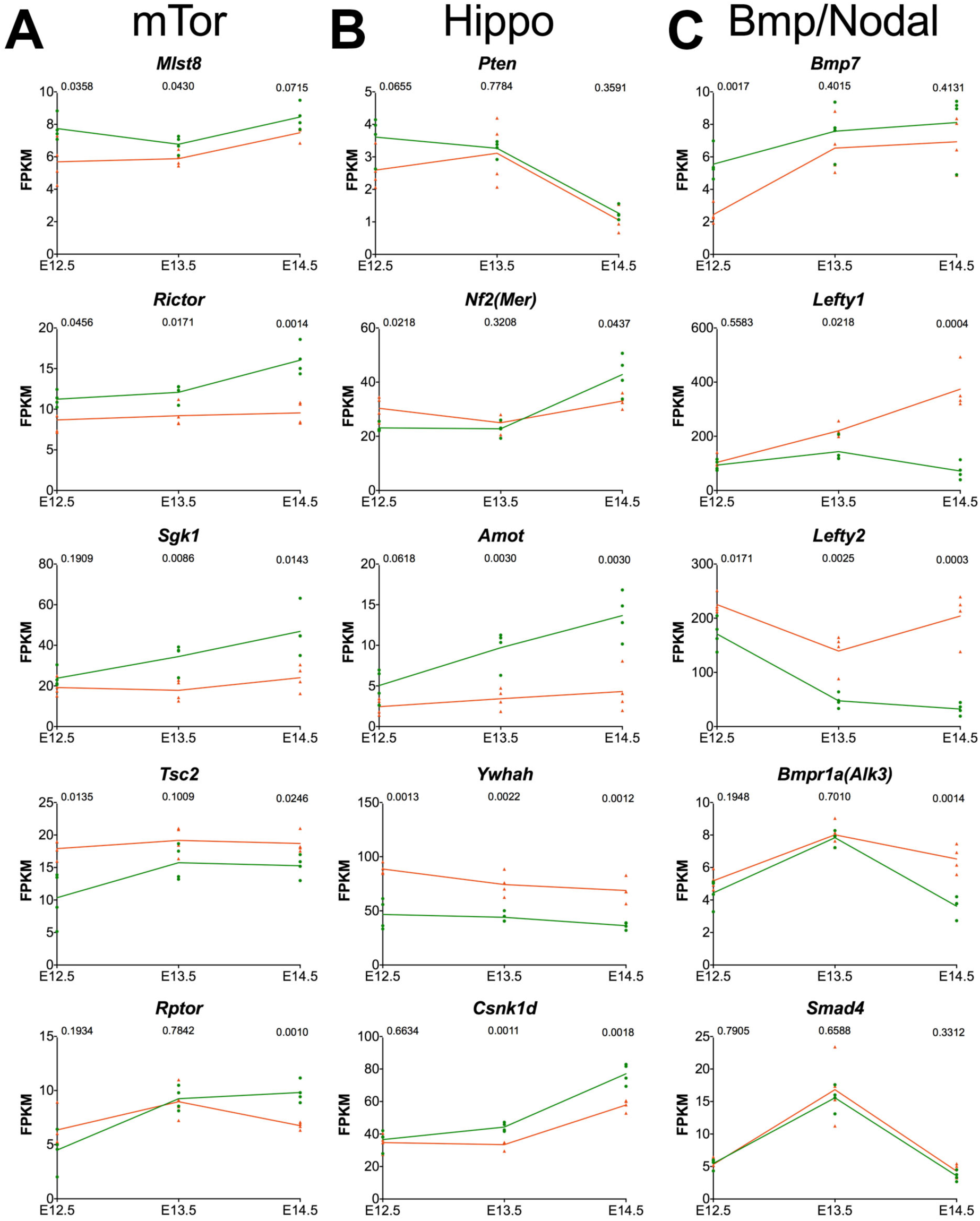
Expression of genes in the mTor, Hippo and Bmp/Nodal pathways are altered in *DND_1_*^*Ter*/*Ter*^ mutants. Expression of other members of the **(A)** mTor, **(B)** Hippo and **(C)** Bmp/Nodal pathways from Fig. 5 is misregulated in mutants with some mapping as transcript targets of DND_1_ (Fig. 5A).

